# Spatial epigenomic niches underlie glioblastoma cell state plasticity

**DOI:** 10.1101/2025.05.09.653178

**Authors:** Sam Kint, Subhi Talal Younes, Shuozhen Bao, Gretchen Long, David Wouters, Erin Stephenson, Xing Lou, Mei Zhong, Di Zhang, Graham Su, Archibald Enninful, Mingyu Yang, Huey-Miin Chen, Katrina Ellestad, Colleen Anderson, Jennifer Moliterno, John Kelly, Jennifer A Chan, Alejandro Sifrim, Jiangbing Zhou, Ana Nikolic, Rong Fan, Marco Gallo

**Affiliations:** Arnie Charbonneau Cancer Institute, Cumming School of Medicine, University of Calgary, Calgary, AB T2N 4N1, Canada; Alberta Children’s Hospital Research Institute, Cumming School of Medicine, University of Calgary, Calgary, AB T2N 4N1, Canada; Department of Pediatrics, Baylor College of Medicine, Houston, TX, 77030, United States of America; Cancer and Hematology Center, Texas Children’s Hospital, Houston, TX, 77030, United States of America; Department of Biomedical Engineering, Yale University, New Haven, CT, 06520, United States of America; Department of Neurosurgery, Yale University, New Haven, CT, 06520, United States of America; Department of Human Genetics, University of Leuven, KU Leuven, Leuven, 3000, Belgium; KU Leuven Institute for Single Cell Omics (LISCO), University of Leuven, KU Leuven, Leuven, 3000, Belgium; KU Leuven Institute for Artificial Intelligence (Leuven.AI), University of Leuven, KU Leuven, Leuven, 3000, Belgium; Department of Pathology and Laboratory Medicine, University of Calgary, Calgary, AB T2N 2T9, Canada; Department of Cell Biology, Yale University, New Haven, CT, 06520, United States of America; Department of Pathology, Yale University School of Medicine, New Haven, CT, 06520, United States of America; Systems Biology Institute, Integrated Science and Technology Center, West Haven, CT, 06516 United States of America; Yale Stem Cell Center and Yale Cancer Center, Yale University School of Medicine, New Haven, CT, 06520, United States of America; Human and Translational Immunology, Yale University, School of Medicine, New Haven, CT, 06520, United States of America; Dan L Duncan Comprehensive Cancer Center, Baylor College of Medicine, Houston, TX 77030, United States of America

**Keywords:** Spatial genomics, spatial ATAC, spatial multiomics, scATANC, glioblastoma, glioma, brain tumors, epigenetics, epigenomics, plasticity, cell state transitions

## Abstract

*IDH*-wildtype glioblastoma (GBM) is an aggressive brain tumor with poor survival and few therapeutic options. Transcriptionally-defined cell states coexist in GBM and occupy defined regions of the tumor. Evidence indicates that GBM cell states are plastic, but it remains unclear if they are determined by the underlying epigenetic state and/or by microenvironmental factors. Here, we present spatially-resolved epigenomic profiling of human GBM tissues that implicate chromatin structure as a key enabler of cell plasticity. We report two epigenetically-defined and spatially-nested tumor niches. Each niche activates short-range molecular signals to maintain its own state and, surprisingly, long-range signals to reinforce the state of the neighboring niche. The position of a cell along this gradient-like system of opposing signals determines its likelihood to be in one state or the other. Our results reveal an intrinsic system for cell plasticity that is encoded in the chromatin profiles of two adjacent niches that dot the topological architecture of GBM in cartesian space.

## BACKGROUND

*IDH*-wild type glioblastoma (GBM) is an aggressive brain tumor with poor survival because of lack of effective therapeutic options^1^. GBM is a malignancy with complex biology and is characterized by extensive intratumoral cellular heterogeneity at the genetic, transcriptional and epigenetic levels^2–5^. In recent years, single-cell genomic approaches^6–9^ have allowed unprecedented precision in characterizing transcriptional heterogeneity and it is now recognized that cells with different transcriptional profiles, or cell states, coexist in the same tumor. Although several nomenclatures have been proposed to name GBM transcriptional cell states, they are all consistent with the presence of two broad categories: Neural/oligodendrocyte progenitor-like (OPC-like) and astrocytic/mesenchymal-like (MES-like) cell states^10^.

Several recent studies have adopted spatial transcriptomic platforms to investigate the distribution of cell states in cartesian space^11–15^. These studies have shown that a subset of GBM specimens display spatial segregation of transcriptionally defined cell states, whereas others are largely unstructured. Local enrichment of cell states has been predicated on the influence of the local microenvironment, especially in terms of neighboring non-neoplastic brain cell types and proximity to regions of marked hypoxia. On the other hand, subclonal genetics appears to have only limited effects on GBM transcriptional states^11^. These conclusions contrast with previous bulk and single-cell genomic analyses^9,16^, which revealed correlations between specific genetic lesions and transcriptional profiles. One may hypothesize that the transcriptional state is shaped largely by epigenetic regulation and influenced by cues from the tissue microenvironment. More studies are therefore needed to spatially map the epigenetic landscape of tumors and disambiguate the intricate relationships between genetic, epigenetic, and transcriptional states of GBM cells.

Recent studies also suggest GBM cell states are plastic. Computational analyses based on single-cell RNA-seq data inferred unidirectional transitions from mesenchymal-like to proneural-like cell states^17^. Further, transcriptionally hybrid cell states have been reported in primary specimens using single-cell RNA-seq^18,19^, further supporting the possibility of dynamic cell state transitions. However, there is a dearth of information on mechanisms and dynamics of cell state transitions (i.e. an individual cell changing state) in human surgical specimens. Identifying mechanisms and principles of cell plasticity in cancer could be of high clinical significance because they are believed to underlie therapy resistance by enabling cells to shift to non-targetable states^17,20^.

We hypothesized that if transcriptional plasticity occurs in GBM, it is likely to depend on permissible chromatin and epigenetic modulation. Assays that profile epigenetic and chromatin landscapes could therefore provide information on mechanisms of transcriptional shifts that enable cell plasticity. We further hypothesize that epigenetic cell state transitions are dependent on the tissue niches and regulated by microenvironmental factors. To test the role of chromatin programs in cell plasticity, we profiled the chromatin accessibility landscapes of GBM surgical specimens with spatial ATAC (assay for transposase-accessible chromatin^21^). This technique generates spatially resolved genome-wide maps of accessible chromatin regions, which correspond to active cis-regulatory elements and actively transcribed genes. Additionally, through inference of transcription factor (TF) motif enrichment in active chromatin regions, spatial ATAC data provide information on potential mechanisms of transcriptional regulation across the cartesian space of a tumor sample. To the best of our knowledge, this is the first spatial epigenomic map of any brain tumor type.

In addition, we generated spatial multiome (ATAC and RNA-seq), multiplex protein imaging, and single-nucleus ATAC data for select samples in our cohort which, upon integration, resulted in characterization of GBM surgical specimens with unprecedented resolution. Our results show that GBM cells are organized in two epigenetically distinct and spatially segregated niches that communicate and reinforce each other’s behavior. Transitions between niches are enabled by asymmetric cell-cell communication, which activates or represses key developmental genes that are found in regions of accessible chromatin in both states. Our results therefore provide a mechanistic framework for GBM cell plasticity that is predicated on the position of an individual cell in fields of opposing signals that emanate from two main epigenetically defined niches.

## RESULTS

### Spatial epigenomic profiling of GBM surgical specimens

Single-cell genomic studies have determined that GBM is characterized by extensive intra-tumoral epigenetic heterogeneity^3,4^. However, it is not known whether cells with different epigenetic profiles are geographically separated or intermixed in a tumor. To address this question, we performed a spatially resolved assay for transposase accessible chromatin (spatial ATAC^21^) on 28 *IDH-*wildtype GBM surgical resections from 24 patients (**Figure 1A**). In 4 tumors, we profiled geographically separated core and peripheral regions (contrasting and non-contrast enhancing, respectively). To complement our findings from spatial epigenomic analyses, we also employed spatial multiome^22^ (n = 4), single nucleus ATAC (n = 3) and multiome (n = 2), PhenoCycler^23,24^ (n = 6), and external datasets^6,11^. Our assembled analyses provide precise ability to decipher epigenomic features of GBM and explore the spatial relationships thereof (**Figure 1A-D**).

**Figure 1.**
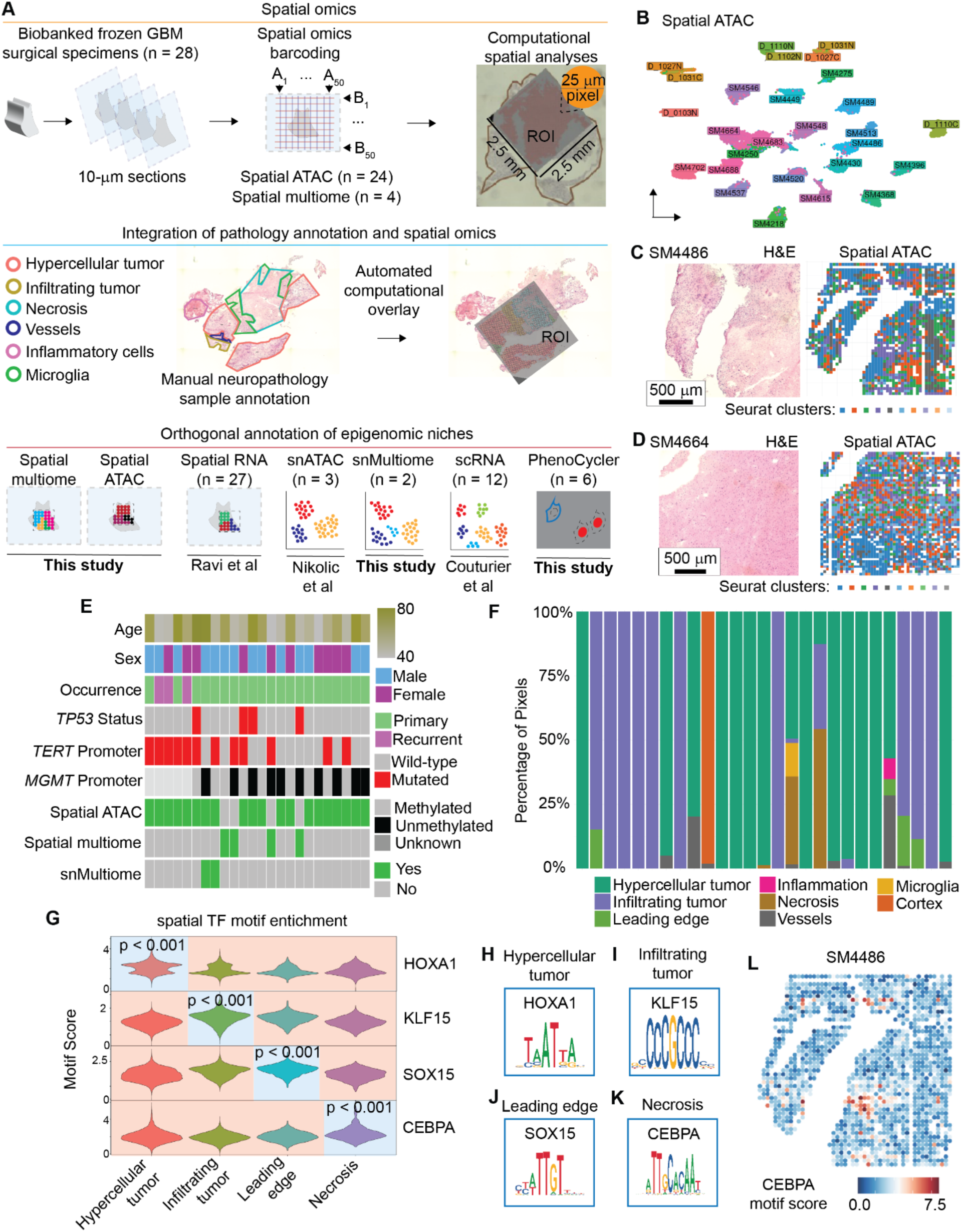
Spatial epigenomic profiling of glioblastoma. (A) Overview of experimental approach for the study, including the multiple datasets used for validation of key findings. (B) UMAP plot of spatial pixels with. Pixels are colored by sample. (C – D) Representative H & E as well as spatial plot of Seurat clusters demonstrating high level structure of the spatial epigenome. (E) Oncoplot showing the major clinical and pathologic characteristics of the patient cohort. (F) Stacked barplot showing percentage of each histologic annotation for each sample. (G) ChromVAR motif scores for TFs of interest across histologic annotations. (H – K) Motifs for each of the major histologic annotations. (L) Representative spatial plots of ChromVAR motif score for CEBPA; note the enrichment in a small area that histologically was determined to contain inflammatory cells (microglia).

Overall, the cohort was representative of typical GBM molecular alterations with respect to *TP53* mutation status, *TERT* promoter mutations, and *MGMT* promoter methylation status (**Figure 1E**). The average age of the patients was 60.7 years (range: 47 – 79 years) with a 1.4:1 male:female ratio, in line with epidemiological trends for this tumor type^25,26^. Patient information is summarized in **Table S1**. Our spatial epigenomic platform (AtlasXomics) achieved a resolution of 25 µm per pixel. In total, we generated data for 54,000 pixels across all samples, with 31,863 passing stringent quality control criteria (see Methods and **Figure S1A-E**).

For each sample, the section immediately adjacent to the one profiled by spatial omic technology was used for hematoxylin and eosin (H&E) staining and manually annotated by a neuropathologist according to clinical parameters relevant to the pathobiology of GBM, including hypercellular tumor, infiltrating regions, areas of necrosis, presence of blood vessels, and microglia and inflammatory cells (**Figure 1A)**. The histologic annotations were then overlayed on the section profiled by spatial ATAC using an automated method (see Methods section)^27^. In 23 samples, we mostly captured one type of histological area, while 5 samples spanned multiple histological classifications (**Figure 1F**). This approach enabled studies of the chromatin states that are enriched in distinct histological regions. For instance, we identified significant differences in TF motif enrichment in accessible chromatin regions across histologic compartments (**Figure 1G**). Developmental-associated TFs (e.g. HOXA1) were enriched in hypercellular tumors (**Figure 1H**). The leading edge and infiltrating portions of the tumor demonstrated enrichment for TFs associated with glial cells or glial cell development, including KLF15 in infiltrating tumor and SOX15 at the leading edge (**Figure 1I–J**). As expected, accessible chromatin regions in necrotic areas of the tumor had enrichment for inflammation associated TFs, including CEBPA **(Figure 1K–L**). Overall, our datasets enabled spatially informed exploration of epigenetic profiles in a clinically relevant GBM cohort.

### Cortical– and white matter-like chromatin modules impart topological organization to GBM

Given the complexity of the spatial organization of the GBM microenvironment and its multiple cell types, we first set out to identify the key cellular and transcriptional features of our dataset using an unsupervised approach. Given the robustness of TF motif profiles and their importance in specifying cell type^28^, we elected to use ChromVAR motif scores for this approach. We computed global motif deviations deviations in pixels with high global similarity scores using neighbourhood clustering to generate pseudopixels and subsequently ran Weighted Gene Network Correlation Analysis (WGCNA)^29^ on the clustered motif deviations (**Figure 2A**). Our motif WGCNA identified 10 distinct modules. Based upon manual review of the genes within those modules and correlation of module scores with our previously annotated single-nucleus ATAC-seq dataset^30^ (**Figure S2A-E**), we then assigned 8 out of 10 modules a descriptor (**Figure 2B** and **Figure S2E-L**). The two most prominent modules were (1) a cortical module, associated with motifs such as ATOH1, SNAI1 and ZEB, the latter having been associated with reactive astrocytosis^31^, and (2) a white matter module associated with transcription factors associated with myelination including ATF3 and CREB^32^. Another module was associated with mature neural and glial cells and included proneural transcription factors and mature oligodendroglial regulators such as NEUROD1, NEUROG1, OLIG1, and OLIG2. We identified three tumor-associated modules, including an ASCL1/neural progenitor-like module a HOX-like module enriched for transcription factors such as GSX1/2, HOXB3, and MEOX2, and a module enriched for the AP-1 transcription factor family, including FOS and JUN family members. Lastly, we noted two microenvironmental modules associated with mature myeloid cells (enriched for SPI1 and the ETV family) and another associated with mature oligodendrocytes. These modules had distinct geographic distributions, as can be demonstrated in sample D_1110C, with the cortical and white matter modules being anticorrelated (**Figure 2C** – **E**) as well as the dual ASCL1/NPC and AP-1 tumor modules (**Figure 2F** – **G**). Of note, the AP-1 module was frequently associated with perinecrotic zones, in keeping with the known role of these transcription factors in regulating the hypoxia response^33,34^. We observed clear zonation of these states (**Figure 2G**), with enrichment of AP-1 adjacent to the necrotic region, enrichment of the myeloid module in a focus of inflammatory cells, and enrichment of the ASCL1/NPC-like state further from the areas of necrosis and inflammation. These modules not only captured some of the histologic features of the samples, but also, in the case of the AP-1/hypoxia module, highlighted distinct programs not apparent with H&E staining.

**Figure 2.**
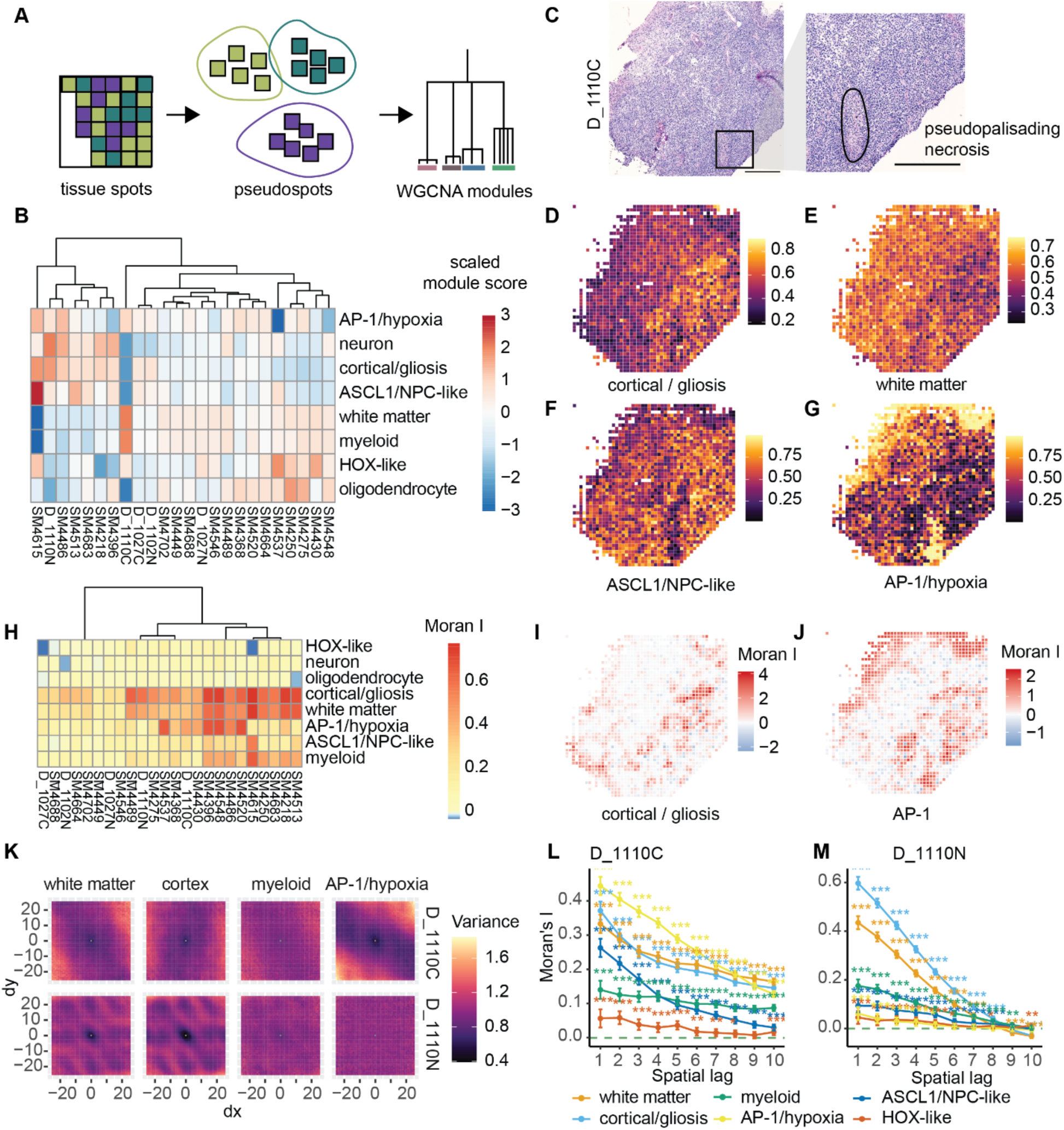
Cortical and white matter epigenetic modules provide scaffolds for GBM organization. (A) Overview of WGCNA strategy. (B) Scaled motif module scores across different samples. (C) Representative H&E image. (D – G) Distributions of module scores for the cortical / gliosis (D), white matter (E), ASCL1/NPC-like (F) and AP-1/hypoxia (G) modules. (H) calculated Moran’s I scores for motif module scores. (I – J) Representative module Moran’s I scores for the cortical/gliosis (I) and AP-1 modules (J). (K) Spatial variogram maps of select motif modules in paired samples core (D_1110C) and tumor periphery (D_1110N). (L – M) Spatial correlograms of select motif modules in paired samples tumor core and periphery.

We next sought to further classify to what extent these modules were associated with local structure in our samples by computing spatial autocorrelation of our module scores using Moran’s I^35^ (**Figure 2H**). The cortical and white matter modules showed high autocorrelation in most samples, consistent with broad baseline domains of underlying cortex and white matter. In contrast, the AP-1/hypoxia signature was variable, showing high autocorrelation in a subset of samples, and relatively minimal structure in others, highlighting that this pattern is only spatially organized in a subset of samples. The neuron, oligodendrocyte, and HOX-like clusters showed much lower autocorrelation, consistent with association with single or infiltrating cells, and the myeloid and ASCL1 related programs were intermediate, showing some structure in a subset of cases. Calculation of local spot Moran I for the highly autocorrelated programs showed that their structure varied within samples and was composed of smaller local domains of enrichment interspersed with regions of exclusion (**Figure 2I-J**). We then explored this in more detail in a set of spatially segregated samples from the same tumor, D_1110N and D_1110C, representing an infiltration zone (N) and tumor core (C), respectively. Spatial variogram maps of select modules showed high structure in both samples for cortical and white matter programs, but strong structure for the AP-1 and myeloid programs only within the tumor core and not the infiltration zone (**Figure 2K**). Examination of spatial correlograms, which indicate decay of autocorrelation over distance and approximate domain size, revealed a similar pattern, with high structure of the cortical and white matter programs in both samples, but striking enrichments in autocorrelation of both the AP-1/hypoxia and ASCL-1/proneural programs (**Figure 2L-M**). This is consistent with a model in which cortical and white matter structures are important organizing principles within all regions of the tumor, while hypoxia is a more significant driver of tumor phenotype within the tumor core.

### Genetic subclones defined by extrachromosomal amplifications are spatially segregated in GBM

To better understand the spatial epigenetic variability in GBM, we identified the top 1000 genes with highly variable gene activity score across our dataset. Because we were interested in which genes may be important in forming broad spatial domains (regions of the tumor with similar epigenetic profiles), we calculated spatial autocorrelation using Moran’s I for each of these genes. Clustering of the top 100 spatially variable genes based upon that autocorrelation score identified six distinct clusters. Interestingly, four of these clusters consisted entirely of genes classically associated with genomic amplifications in GBM (**Figure S3A**). For example, cluster 6 consisted of the genes *EGFR*, *ELDR*, and *LANCL2* which are in proximity to one another on chromosome 7 and are frequently co-amplified in GBM. Additional clusters of interest include one associated with putative *MDM2* amplification (cluster 5), and *CDK4* amplification (cluster 4), both of which are located on chromosome 12. The remaining clusters of genes with similar autocorrelation patterns likely represent normal cortex, white matter, and myeloid cells as those clusters contained genes such as *ROBO2* and *SOX8*, respectively.

Thus, pixels which exhibit amplification of *EGFR*, *MDM2*, or *CDK4* are more often adjacent to one another than would be anticipated by chance (i.e. they exhibit high autocorrelation). Since such amplifications are often subclonal in GBM^36^, this data suggests that genetic subclones may spatially co-localize within tumors.

To visualize these putative gene amplifications, we computed scores consisting of the sum of gene activities for all genes in each cluster and scored each pixel. Stratifying spots based on scores for an autocorrelation cluster associated with non-malignant genes (cluster 2) and a cluster associated with *EGFR* pathologic gene amplification (cluster 6) was able to distinguish those spots which likely contain entirely normal tissue, mixed normal and tumor cells, and pure tumor (**Figure 3A**). Samples with high tumor cellularity contained broad sheets of pixels with high scores for putative gene amplifications (**Figure 3B-C**). In contrast, in areas of infiltrating tumor and low tumor cellularity, *EGFR* amplification highlighted individual pixels of tumor infiltration (**Figure 3D– E** and **Figure S3B–E**). Neighborhood analysis of first, second, and third order pixel neighbors showed relative homogeneity of *EGFR* amplification in areas of hypercellular tumor with substantially lower neighborhood scores for *EGFR* amplification at the leading edge (**Figure 3F-G**). We next built neighborhood graphs of *EGFR*-amplified domains across all samples with this amplification in our cohort. We then quantified domain sizes, demonstrating that most of our samples contained multiple small, disconnected *EGFR* amplification domains (**Figure 3H-I**). These data indicate that *EGFR*-amplified tumor cells locally segregate together within the tumor.

**Figure 3.**
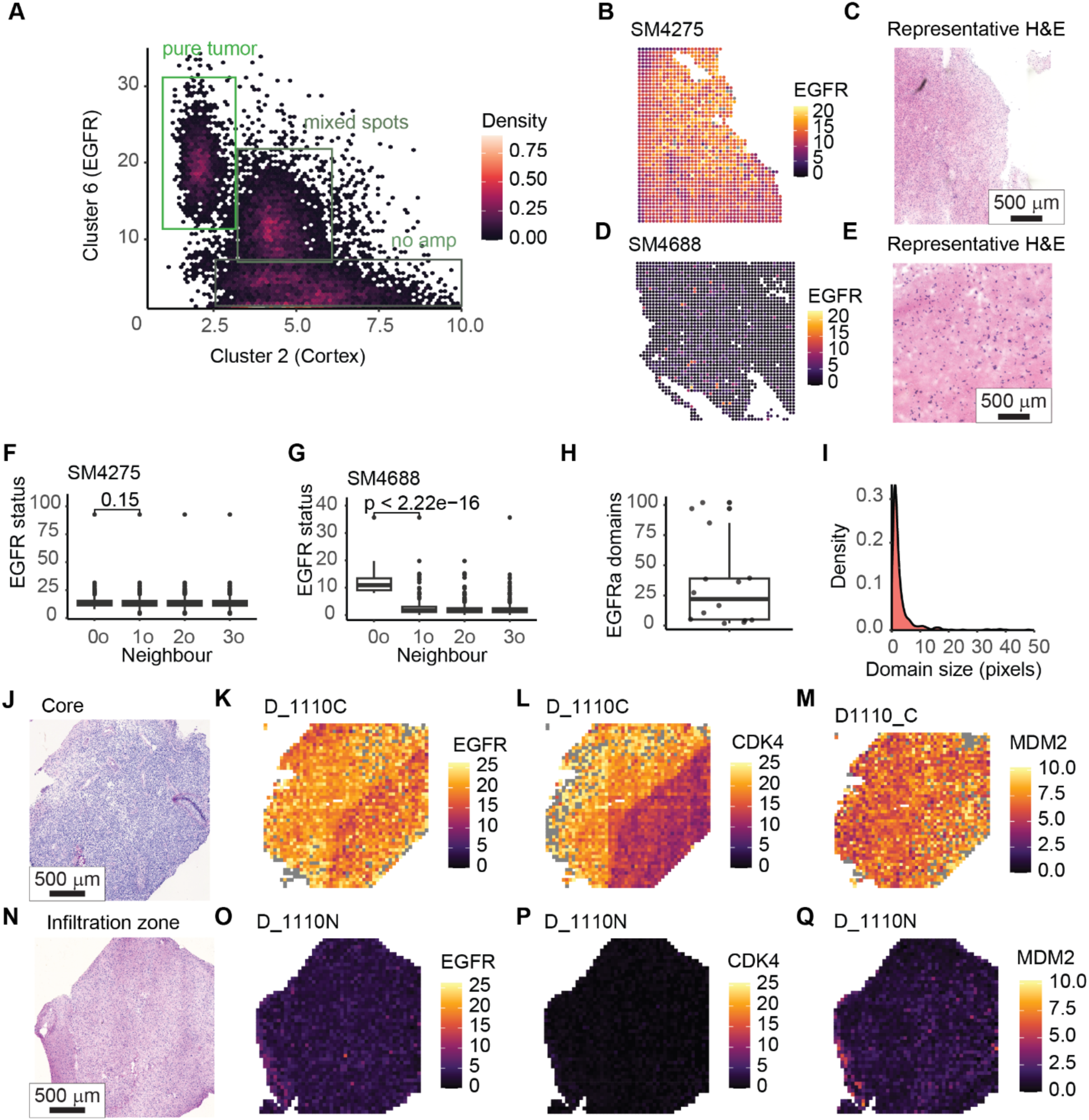
Genetic subclones are spatially localized in topological space. (A) Density plot of EGFR amplification versus cortex score across all samples. (B – E) Representative *EGFR* amplification scores and H&E images of a high and low-cellularity EGFR-amplified sample. (F – G) Neighborhood *EGFR* scores in two *EGFR*-amplified samples. (H – I) Summary of *EGFR* amplification domains per sample and domain sizes. (J – Q) Spatial plots of indicated gene cluster for a matched tumor core and peripheral sample as well as representative H & E of those samples.

We next leveraged one of our multi-regionally mapped samples (D_1110) to examin differences in subclonal composition between the core and periphery of the tumor. Amplifications of *EGFR* and *CDK4* were present in the core of the tumor, yet sparse and focal in the peripheral sample despite high tumor cellularity (**Figure 3J-Q**). Furthermore, *MDM2* amplification remained present in the peripheral, infiltrating zone of the tumor (**Figure 3Q**). These data support gain of multiple subclonal chromosomal rearrangements within the tumor core, suggesting that there is ongoing tumor evolution and expansion within the core. Similar patterns of spatially segregated genetic subclones were seen in other samples. For example, sample SM4250 showed spatially segregated regions of *MDM2* and *EGFR* amplification (**Figure S3F–I**). These subclonal amplifications were confirmed in single nucleus ATAC-seq from the same sample (**Figure S3J–L**). Similarly, sample SM4702 exhibited prominent CDK4 amplification but only small areas of MDM2 amplification (**Figure S3M-O**),

Taken together, these data show that subclones of GBM cells – here distinguished by chromosomal amplifications – exhibit spatially distinct localizations both within core regions of the tumor and can show differences between the core and periphery within same tumor.

### GBM harbors spatially nested cellular niches

To better understand the spatial organization of GBM chromatin states, we utilized NicheCompass^37^, a graph neural network-based computational tool that enables the identification of putative cellular niches (**Figure 4A**). In a partially supervised manner (i.e. informed by known gene programs), the neural network assigns each spot a score for latent gene programs. Subsequent clustering on the latent gene program space then produces a spatial map of cellular niches. NicheCompass relies primarily on spatial RNA data; thus, we used the 4 samples that we profiled with spatial multiome for this analysis. One sample was excluded due to low RNA quality.

**Figure 4.**
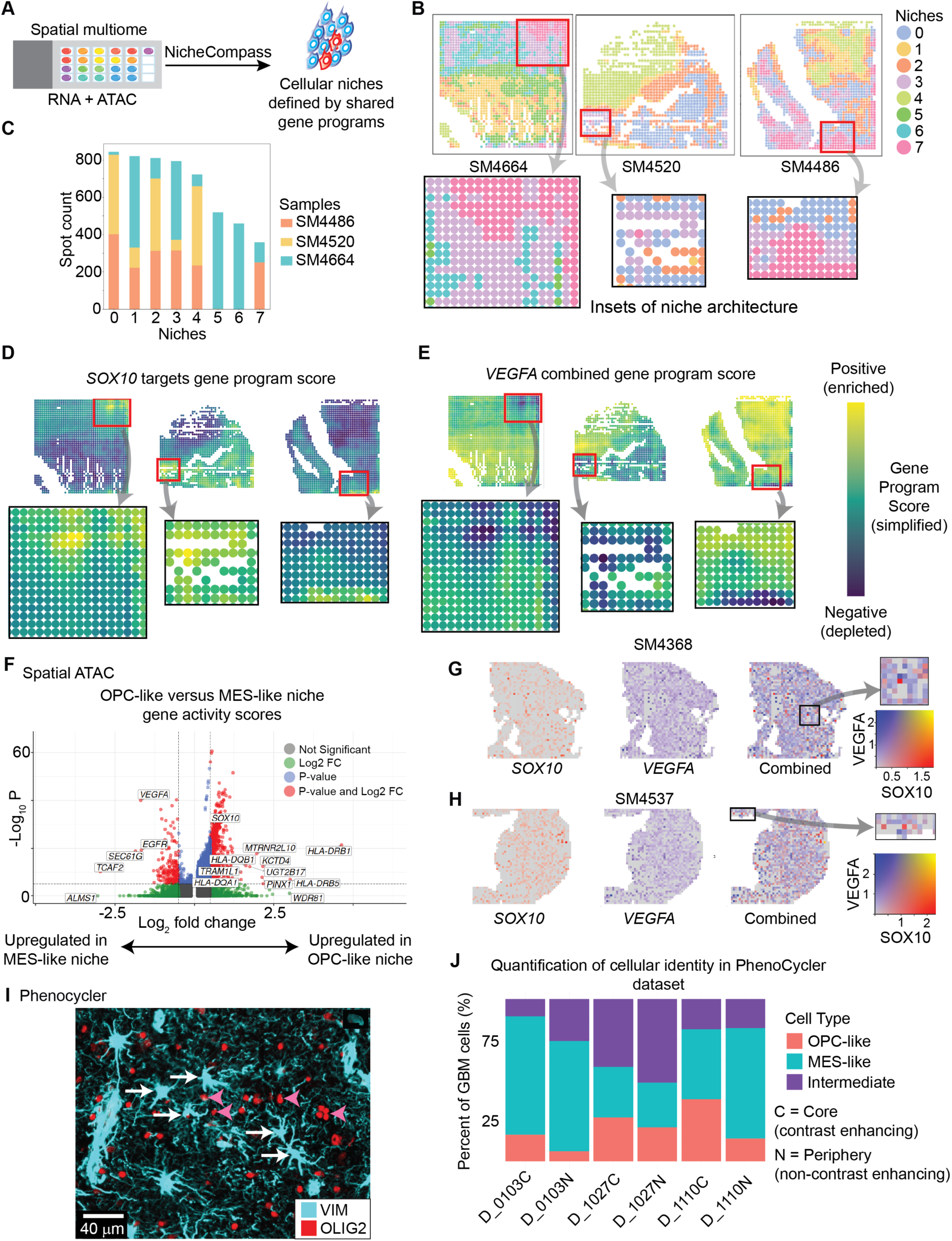
Spatially-segregated epigenetic niches in GBM. (A) Overview of NicheCompass workflow. (B) Spatial plots of NicheCompass-defined cellular niches with regions of interest highlighted. (D) Spatial plot of *SOX10* targets gene program across all samples. (E) Spatial plot of *VEGFA* combined gene program score. (F) Volcano plot of gene activity scores for pixels belonging to niches 3 and 7 (OPC-like) versus niche 0 (MES-like). (G – H) Representative spatial plots of SOX10 and VEGFA gene activity scores highlighting regions of OPC-like niches surrounded by MES-like niches (I) Representative microscopy image from PhenoCycler data showing the presence of MES-like (white arrows) and OPC-like cells (magenta arrowheads) within the core of a GBM tumor. (J) Quantification of cellular identity in PhenoCycler data. MES is defined as VIM positive, OPC as OLIG2 positive, and intermediate as positive for both markers.

With this approach, 8 primary cellular niches were identified (**Figure 4B**). 6 of the 8 niches were present across all 3 samples (**Figure 4C**). We noticed that some niches exhibited an intriguing spatially nested architecture (insets in **Figure 4B**). Analysis of differential gene programs between the inner and outer niches determined that *SOX10-* and *SOX2*-associated gene programs were enriched in the inner niche (**Figure 4D** and **Table S2**; log Bayes factor 9.0 and 5.3, respectively). On the other hand, *VEGFA-* and *CD44-*associated gene programs were relatively depleted within these niches (**Figure 4E** and **Table S2;** log Bayes factor –4.7 and –2.4, respectively). Additional gene programs with differential enrichment included *ANGPTL4, BMP4,* and *WNT4* (see **Table S2;** log Bayes factor 27, 12.5, and –4, respectively).

To better characterize the inner and outer cellular niches in GBM, we determined differential gene activities between the inner niches and their immediate surrounding niche using spatial ATAC data. We identified several known GBM-associated genes with significant differential gene activity scores between the inner and outer niches (**Figure 4F**), including *SOX10* within the inner niche and *VEGFA* and *EGFR* in the outer niche. These results suggest an OPC-like profile for the inner niche and a MES-like profile for the outer niche.

To determine whether this architecture of inner and outer niches was present across our epigenomic dataset, we reviewed the differential gene activities to identify potential markers for each cell identity. This curation nominated *SOX10* as a candidate marker for the inner niche and *VEGFA* for the outer niche. Using gene activity cutoffs of 0.5 for *SOX10* and *VEGFA*, we found that 13% of pixels were positive for *SOX10*, 22% were positive for *VEGFA*, 7% were positive for both, and 58% were negative for both (**Figure S4A)**. The high percentage of pixels negative for both *SOX10* and *VEGFA* is likely due to admixture of non-neoplastic and infiltrating tumor cells (see **Figure 1C**).

With this assignment system based on *SOX10* and *VEGFA* gene activity, 27 of 28 samples in our cohort showed spatial architecture characterized by small nests of *SOX10*-positive pixels surrounded by sheets of *VEGFA*-positive pixels (**Figure 4I**-**J** and **Figure S4B**). These data suggest that even OPC-like and MES-like GBM cells co-exist in spatially segregated niches both in the core and at the periphery of the tumor, and not exclusively at the periphery as was previously thought^15,20,38,39^.

To further validate whether OPC-like and MES-like cells indeed coexist within the same regions of tumor, we interrogated GBM cell states at the protein level using PhenoCycler^23,24^, a microscopy platform which allows interrogation of multiple protein at the single cell and subcellular level. However, in our study, even in samples from the core of the tumor, we identified OPC-like and MES-like cells (here demarcated using OLIG2 and Vimentin) in adjacent niches (**Figure 4H**). Furthermore, in paired core and peripheral samples from the same tumor, we readily identified OPC-like and MES-like cells in both the core and periphery of each tumor (**Figure 4I**); indeed, by percentage each tumor had more OPC-like cells in the core. Finally, in a previously published spatial transcriptomic dataset of GBM specimens^11^, we again identified nests of *SOX10*-positive pixels surrounded by broad sheets of *VEGFA*-positive pixels in 13 of 27 cases (**Figure S5**), indicating the co-existence and spatial segregation of OPC-like and MES-like cells.

Altogether, results from four orthogonal approaches (spatial multiome, spatial ATAC, spatial transcriptome, and PhenoCycler) across independent cohorts of GBM samples reveal that OPC-like and MES-like cellular niches co-exist as adjacent and spatially segregated niches in both the tumor core and margin.

### Cell-cell communication suggests mechanisms of cellular plasticity in GBM

Given the spatial relationship of OPC-like and MES-like niches, we wondered whether these niches communicated with each other via cell-cell signaling pathways. Therefore, we first used NicheCompass to identify signaling programs in spatial multiome samples (**Figure 5A**). *PDGFB* signaling was prominent within the OPC-like niche (niches 3 and 7) (**Figure 5B**). Since PDGFB specifies OPCs during normal brain development and promotes their rapid division^40,41^, these data suggest that OPC-like GBM cells maintain their own cell state and promote self-proliferation, at least in part, by autocrine and paracrine PDGFB signaling.

**Figure 5.**
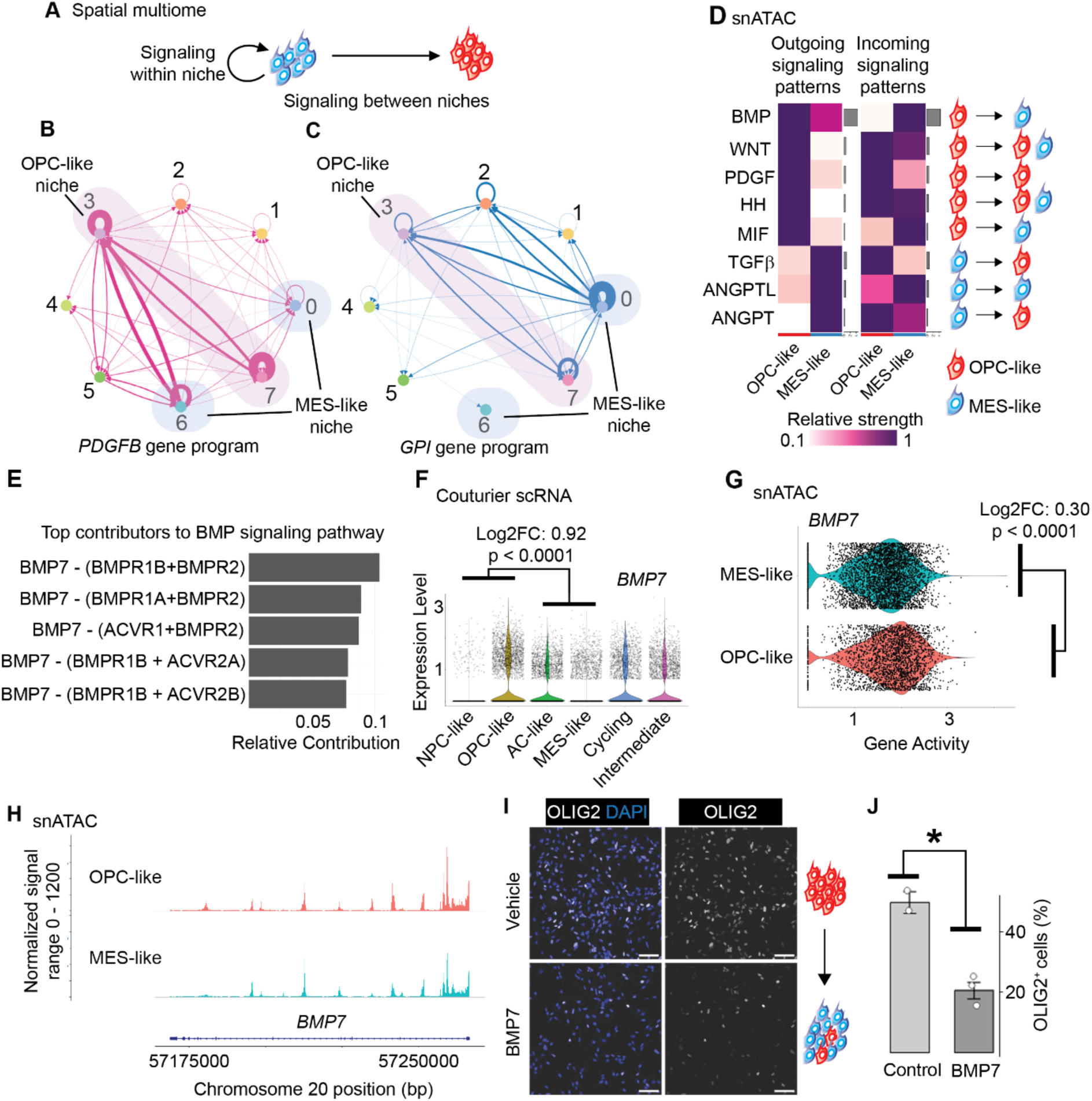
GBM niches communicate to reinforce each other’s behavior. (A) Schematic demonstrating how data are portrayed for panels B and C. (B – C) Signaling programs as defined by NicheCompass demonstrating PDGFB signaling within the OPC-like niche and GPI signaling from the MES-like niche to the OPC-like niche. (D) Heatmap of signaling probabilities as defined by CellChat based on snATAC data of malignant GBM cells. (E) Top ligand-receptor pairs contributing to BMP signaling pathway. (F) Violin plots of *BMP7* expression in single cell RNA sequencing data from Couturier and colleagues. (G) Violin plot of gene activity score from snATAC dataset. Note the relatively modest difference in gene activity. (H) Coverage plot of the *BMP7* gene. Note relatively similar chromatin landscape. (I) Fluorescence microscopy images of GBM cell culture (GSC016) under control conditions and after treatment for 72 hours with recombinant BMP7 (n = 2 – 4 wells per condition). Scale bar indicates 100 µm. (J) Quantification of percent OLIG2 positive cells in (I).

On the other hand, a signaling program associated with *GPI* was prominent in the MES-like niches (niches 0 and 6) and was directed to the inner niches as well as having a prominent self-signaling nature (**Figure 5C**). The OPC-like and MES-like niches therefore have their own signaling programs that may enable inter-niche crosstalk.

Orthogonally, we used CellChat^42^ to examine cell-cell crosstalk in our snATAC dataset. Given the greater depth and higher resolution of single nucleus sequencing compared to spatial sequencing, we identified a higher variety of signaling pathways with statistically significant activity between OPC-like and MES-like cells (**Figure 5D** and **Figure S6A**).

We observed two main patterns of signaling: Those within cell types (i.e. the same cell type was both the sender and receiver of the signal) and those between cell types (one cell type was the sender and the other the receiver). We termed these patterns short– and long-range signaling, respectively.

OPC-like cells exhibit short-range signaling of several pathways that maintain or specify OPCs during early development^43^, including Hedgehog, WNT, and PDGF signaling, suggesting that OPC-like cells maintain their own cell state via OPC developmental pathways.

We identified BMP as an example of long-range signaling from OPC-like to MES-like cells, with BMP7 being the predominant ligand and BMPR1B and BMPR2 as the primary receptor pair (**Figure 5E**). The BMP pathway has been previously shown to induce cell state transition in GBM^44^. Long-range signaling from MES-like to OPC-like cells include the ANGPTL and TGF-b pathways (**Figure 5D**), both of which have been shown to promote tumorigenesis and cell migration in GBM^45,46^. These data suggest the presence of a pro-migratory cell signal from MES-like to OPC-like cells. We confirmed cell-type specific production of these key ligands in an independent scRNA-seq dataset, showing that OPC-like cells have higher expression of *BMP7* (**Figure 5F**).

Examination of individual chromatin tracks for signaling molecules involved in the above-described pathways disclosed that rather subtle differences in chromatin accessibility accounted for the observed differences in signaling probability (**Figure 5G–H**). In other words, chromatin exhibited relatively similar accessibility in both cell types at *BMP* loci. A similar pattern of shared open chromatin regions was observed across other highlighted signaling ligands including WNT, PDGF, HH and TGF-b (**Figure S6B – E**). This suggests that differences in expression or signaling probability are likely due to minor alterations at distal regulatory elements.

Furthermore, despite dichotomies in signaling source, both OPC-like and MES-like cells can act as receivers for most pathways examined (**see Figure 5D**), a finding supported again by nearly equivalent chromatin accessibility profiles at receptor loci (**Figure S6F–I**). These data suggest that with relatively minor alterations in chromatin accessibility, both cell types can likely produce and respond to BMP, WNT, PDGF, HH, TGF-b, and ANGPTL signaling.

With this suggestion that GBM cell states are plastic and that niches signal to one another, we wondered whether cell-cell signaling can drive cell state transition. Specifically, we hypothesized that the long range signaling of OPC-like to MES-like cells reinforces the MES-like cellular identity. To test this hypothesis, we treated GBM cells with recombinant BMP7 and evaluated cell type via staining for OLIG2, a marker of OPC-like cells. Treatment with BMP7 dramatically reduced the percentage of OLIG2^+^ cells, suggesting that BMP7 promotes cell state transition to the MES-like state (**Figure 5I–J**).

Taken together, our analysis of cell-cell signaling programs indicates that GBM cells signal both within and between niches and that this signaling can specify GBM cell state. Furthermore, cells are primed to switch signaling ligand production via open chromatin structure at their genomic loci. These data suggests that cell plasticity in GBM is the product of cell signaling pathways that activate or repress downstream transcription at loci that have relatively stable chromatin profiles between cell states.

### OPC-like and MES-like niches have distinct TF programs

To gain more insight into the nature of the OPC-like and MES-like niches, we turned to snATAC sequencing of 5 GBM samples from our cohort, of which three have been previously published^30^. (**Figure 6A**). In total, 10,528 nuclei passed quality control (**Figure S7A-C**). Using canonical cell type markers and Leiden clustering, we were able to distinguish malignant and non-malignant cells (**Figure 6B-C**). Since we were interested in the features of malignant GBM cells, subsequent analyses were performed solely on malignant cell clusters (6,452 nuclei). We verified the ability of *SOX10* gene activity to demarcate the two major cell identities previously described in GBM (**Figure 6D–E** and **Figure S7D – O**).

**Figure 6.**
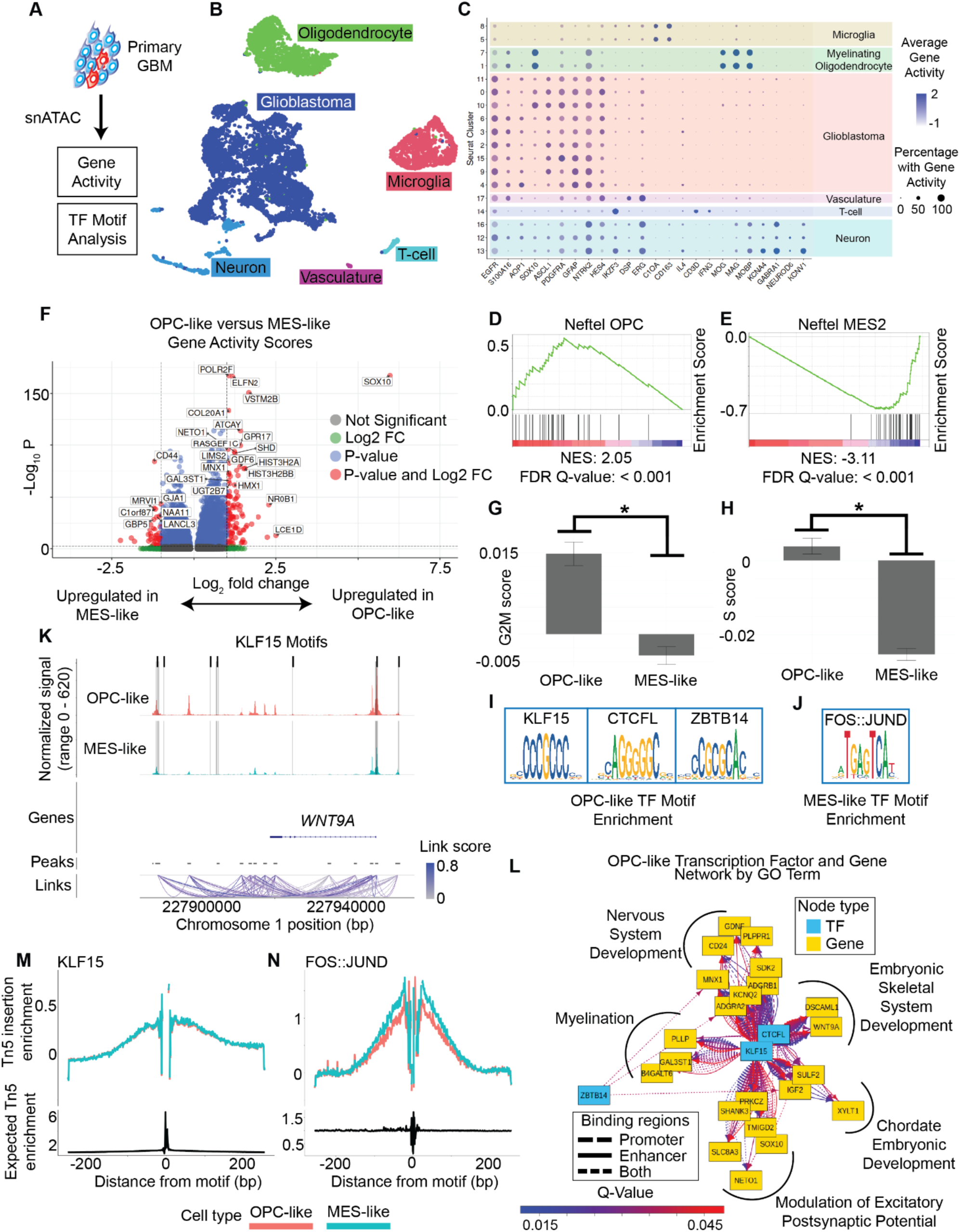
Epigenetic profiles of GBM niches. (A) Overview of experimental design. (B) UMAP plot of snATAC data clustered based off of peaks. Clusters were annotated by gene activity of known cell type markers. (C) Dotplot of gene activities of key cell type markers that formed the basis of cell type calls as shown in panel B. (D – E) Gene set enrichment analysis validating GBM cell type calling. (F) Volcano plot of gene activities comparing OPC-like and MES-like cells. (G – H) Mean and standard error of module scores for G2M and S cell cycle associated genes. (I – J) Motifs of key TFs associated with the indicated cell types. (K) Coverage plot of *WNT9A*, demonstrating relationship of the gene transcription start site to several putative regulatory elements with KLF15 binding sites with strong linking score. (L) Gene regulatory network in OPC-like cells relating TFs to regulatory elements of genes in enriched GO terms. (M – N) TF footprint of indicated TF in OPC-like and MES-like cells.

We next sought to determine the unique chromatin profiles of these two major cell types. Gene ontology analysis of differential gene activities identified GO terms associated with development in OPC-like cells (**Figure 6F** and **Figure S7P**). On the other hand, MES-like cells exhibited enrichment of metabolic-associated GO terms, including those related to glutamate metabolism (**Figure 6F** and **Figure S7Q**). These findings are in line with recent descriptions of the key role metabolism plays in GBM cell state regulation^47^.

We also noted higher gene activity of several cell cycle dependent histone genes in OPC-like cells (**Figure 6F**). Therefore, we sought to further characterize which cell type exhibited higher cycling by comparing module scores for cell cycle-associated genes. We found that whereas each cell type contained cycling cells (**Figure S7R–S**), OPC-like exhibited higher cycling scores than MES-like cells (**Figure 6G-H**; student’s t-test, p-value < 0.0001 for both G2M-phase score and S-phase score), suggesting that OPC-like cells are the predominant mitotic cell type in GBM, a finding which is in agreement with prior studies^9,39^.

Given the key role of TFs in driving cell state, we next examined which TF motifs were enriched in differentially accessible peaks in OPC-like and MES-like cells. We identified significant enrichment of several development-related TFs including KLF15, CTCFL, and ZBTB14 in OPC-like cells (**Figure 6I** and **Figure S7T**), TFs which have been described to be important in key developmental processes^48,49^. TF motif enrichment in MES-like cells was dominated by FOS::JUN (AP-1) TFs (**Figure 6J** and **Figure S7U**).

We next sought to more fully characterize the gene programs being regulated by the transcription factors that were defining these cell identities. Using the GeneHancer database^50^, we searched promoter and enhancer regions of the top GO-term associated genes for enrichment of TF motifs that were also enriched in open chromatin. Doing so identified KLF15 and CTCFL as key TFs, along with ZBTB14 to a minor extent associated with the gene regulatory network in OPC-like (**Figure 6K-L**). A similar analysis failed to identify statistically significant enrichment of specific TF motifs in MES-like cells, likely reflecting a higher degree of heterogeneity in this cell population.

Interestingly, whereas differential motif enrichment does exist between KLF15 and FOS::JUND in OPC-like versus MES-like cells, visualization of the TF footprint demonstrated that each of the key cell-type defining TFs (KLF15 and FOS::JUND) are likely present and bound to DNA in both OPC-like and MES-like cells at similar global levels (**Figure 6M-N**), suggesting that the driver of these cell identities is not differential TF expression or global DNA occupancy of these TFs but rather different genomic targets (loci).

Taken together, these data suggest that the two chief cell types in GBM are characterized by distinct TF motif enrichment and gene programs; however, each respective TF is expressed and bound to DNA in both cell types.

### MES-like states can transition to OPC-like states in GBM

The data shown above indicate that OPC-like and MES-like cells are relatively plastic, as cell type defining TFs are expressed and bound to DNA in both cell types. Therefore, we sought to more fully understand the nature of the relationship of these two cell types in GBM. We applied RNA velocity methods as well as vector field and dynamical systems theory^51^. Here, we used available single-cell RNA sequencing datasets of primary surgical specimens from Couturier and colleagues^6^; in total, 19,410 malignant cells across 12 samples were used for the analysis. Cells were assigned to either MES, AC, OPC, NPC, cycling, or intermediate cell state (not definitively belonging to any of the above five cell states) based on expression of established cell type markers^9^ coupled with Leiden clustering (**Figure S8A – 8G**).

Vector field learning was successful (**Figure S8H**), indicating good performance of the machine learning based algorithm. The vector field recapitulated known relationships between GBM cell types, with MES and AC cells clustering closely together while OPC and NPC cells clustering in proximity of each other and resembling *SOX10*-expressing cells. MES cells are the origin of velocity vectors that point toward all other cell types (**Figure 7A; Supplementary Animation 1**), suggesting that MES cells give rise to all other cell types in GBM. This is further supported by visualization of the Hodge decomposition (a measure of pseudotime), as MES cells exhibit the earliest pseudotime while NPC and OPC cells exhibit the latest (**Figure S8I**–**8J**, **Supplementary Animation 2**).

**Figure 7.**
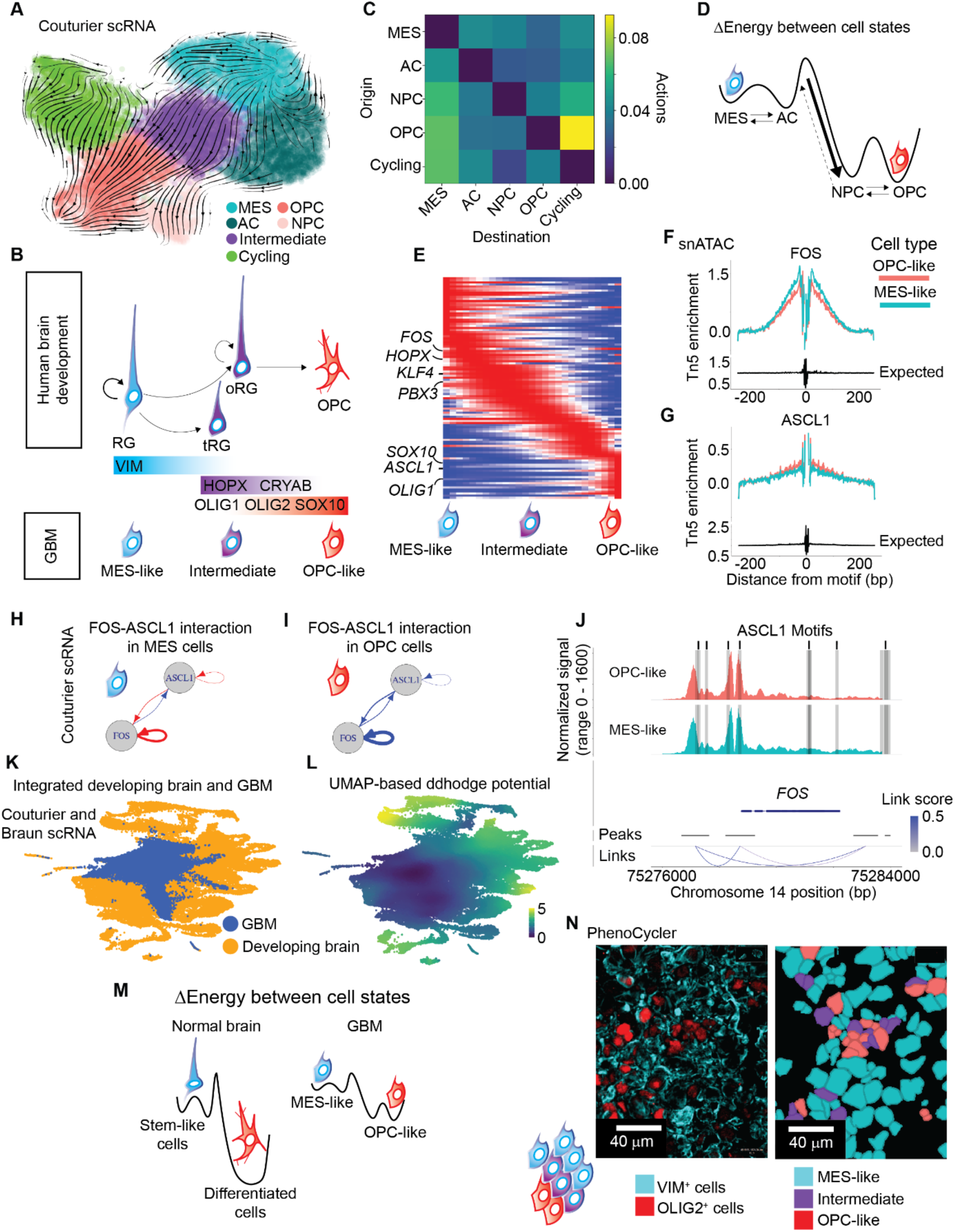
Epigenetically-defined niches provide mechanism for cell plasticity in GBM. (A) Streamline plot of RNA velocity vectors. Note the directionality indicating MES cells as the source of the velocity vectors toward AC, OPC/NPC cell types. (B) Heatmap indicated calculated actions for all cell type transitions. (C) Visual illustration of Hodge potential in GBM, which can be conceived in terms of a Waddington landscape. (D) Illustration comparing normal brain development to the GBM cell relationships. (E) Heatmap of expression for genes with high mean square displacement along MES to OPC transition. TFs of interest are highlighted. (F – G) TF footprints for FOS and ASCL1 from our snATAC data. (H – I) Network graph of cell-type specific relationship between *FOS* and *ASCL1*. Red indicates upregulation while blue indicates downregulation (inhibition). (J) Coverage plot of *FOS*. Note ASCL1 binding sites and the relative similarity of chromatin accessibility across cell types. (K) UMAP plot of integrated GBM and developing brain datasets. Neighbor graph for UMAP embedding is calculated on CCA vectors. (L) UMAP plot colored by ddhodge potential. (M) Illustration indicating the energy landscape of cell state transition in the developing brain versus GBM, noting that cell state transition in GBM requires far less energy compared to the normal developing brain. (N) Representative PhenoCycler microscopy images demonstrating intermediate cell states within the core of a GBM tumor.

If this is the case, given the known parallels between GBM cell states and the normal developing brain, we hypothesized that MES/AC cells would exhibit higher expression of marker genes of radial glia^52^, the primary stem cell of the developing brain^48,53–55^. In support of this hypothesis, MES/AC cells exhibit higher expression of *VIM*, a marker of all radial glia populations (Log2FC 3.06, p-value < 0.001, **Figure S8K**). Beginning at gestational week 17, radial glial populations diverge into the truncated radial glia and outer radial glia populations^48,56^. The gene *CRYAB* marks the former while *HOPX* is enriched in the latter^56^. We observed higher expression of both markers in MES/AC cells as compared to OPC/NPC cells (**Figure S8L** and **8M**); however, the intermediate cell population expressed the highest level of *HOPX* (**Figure S8M**). Since outer radial glia cells are more differentiated than early radial glial cells and OPC cells arise even later in normal development^48,49^, these data suggest that the GBM cell state transition from MES-like to OPC-like recapitulates this developmental paradigm with gene expression changes that mirror the radial glia to OPC differentiation (**Figure 7B**).

To more fully discern the nature of cell state transitions among GBM cells, we computed the least action path between cell states (**Figure S8N**). Here, an action is defined as a movement in the vector field which is not perfectly orthogonal to the velocity vector at that point. Consistent with prior reports, MES and AC cells are more closely related to each other than to OPC and NPC cells (and indeed vice versa), as shown by the need to enter an intermediate cell state to transition from MES to OPC/NPC whereas a MES cell can directly transition to AC (**Figure S8N**). MES and AC cells require the least actions to transition to other cell states while OPC and NPC cells require more actions to undergo transition to other cell states (**Figure 7C**). Furthermore, OPC cells are the destination of most cell state transitions. Taken together, these data show that MES cell states have a higher plasticity potential than OPC cell states (**Figure 7D**).

To more fully characterize the genes which may be driving this partial differentiation, we calculated the mean square displacement (highest change in expression) of genes as cells progressed through pseudotime along the MES to OPC trajectory. Since TFs are key determinants of cell type, we sought to identify which TFs exhibit high mean square displacement. We identified three waves of transcriptional changes with dramatic changes in expression of several important TFs (**Figure 7E**), including *FOS, HOPX, KLF4, PBX3, ASCL1*, *SOX10*, and *OLIG1*. These results strongly imply the requirement of ASCL1 and SOX10 to achieve OPC-like states, in agreement with*SOX10* gene activity being strongly enriched in the OPC-like niches we identified above.

We hypothesized that these TFs are the primary drivers of cell phenotype via their respective gene regulatory networks. The first derivative of the vector field is the Jacobian, an equation which describes gene-by-gene-by-cell interactions^51^. Thus, this calculation produces a putative gene regulatory network. We examined the top positively and negatively related gene regulatory interactions in MES and OPC cells for *FOS* and *ASCL1*. *FOS* upregulates markers of radial glia, including *CRYAB*, as well as the stress-related gene *CLU* (**Figure S8O**). In OPC cells, *ASCL1* downregulates *GAP43* (**Figure S8P**), a key gene involved in microtube formation by MES-like GBM cells in the tumor core, suggesting a mechanism for the previously reported phenomenon whereby OPC cells are unconnected from neighboring GBM cells^38,39^. The activity of these transcription factors was confirmed by inspection of their chromatin footprint in our snATAC data (**Figure 7F – G**). This analysis again showed that whereas there are differences in the TF footprint between major cell types, putative cell-type defining TFs remain present and bound to DNA across cell types, suggesting that subtle alterations in expression and DNA-binding of the TF affect large changes in cell state.

Given the central role of *FOS* and *ASCL1* in cell type determination in GBM, we were intrigued by their interaction in the Jacobian which showed a cell-type specific regulatory interaction. Namely, in MES cells, *FOS* exhibits self-activation while inhibiting expression of *ASCL1* (**Figure 7H**). However, in OPC cells, *FOS* and *ASCL1* exhibit mutual and self-inhibition (**Figure 7I**), highlighting cell-type and context-dependent behavior of these TFs. We sought to validate this putative interaction by motif analysis. We searched known regulatory elements of *FOS* and *ASCL1* for each respective motif. We identified enrichment of *ASCL1* motifs within regulatory elements of the *FOS* gene (**Figure 7J**, p-value <0.01 and FDR q-value < 0.25). No *FOS* binding sites were found within the regulatory regions of *ASCL1* (FDR q-value > 0.5 for all sites interrogated), suggesting that *FOS* indirectly regulates *ASCL1* rather than by direct transcriptional repression. Similarly to findings of the cell-cell signaling molecules, we again noted relatively open chromatin at the *FOS* locus across OPC-like and MES-like cell types (**Figure 7J**).

Taken together, these data support a role for key transcription factors, most notably *FOS* and *ASCL1*, in driving and maintaining cell states within GBM via multiple signaling pathways. Suggestions of high cell plasticity as evidenced by open chromatin at key genomic loci (e.g. *FOS*) across cell states was again noted.

### Cell state transitions in GBM are enabled by a shallow chromatin architecture hierarchy

Whereas our data suggest that there exists a Waddington landscape of partial differentiation in GBM with MES/AC cells at the peak (stem) and OPC/NPC in the valley (differentiated), the comparative heights that may allow cell state transitions (activation energy) can only be described in relative terms. Thus, a key question remains: Is this differentiation valley too steep to allow OPC/NPC cells to “climb” back up the hierarchy or is the valley shallow enough to allow bidirectional cell state transitions? To answer this question, we asked if the transition between GBM cell states has a similar Waddington magnitude to that of normal radial glia to terminally differentiated neurons. We integrated the scRNA-seq dataset of GBM with an scRNA-seq dataset of the developing human brain^57^. This allowed for direct comparison between relative differentiation hierarchies. To conduct such an analysis, our model must first assume that at least some cell states are shared between GBM and the developing brain. This is an assumption well supported by the literature in GBM^8,9,38,52^. We then integrated the two datasets using anchor-based canonical correlation analysis.

GBM cells integrated well with cells of the developing brain (**Figure 7K**). MES, AC, and OPC GBM cells remained closely clustered together whereas NPC cells clustered close to differentiating neuronal lineages in the developing brain (**Figure S9A-B**). We then calculated the vector field on this integrated dataset, substituting the anchoring vectors for the principal components (the latter being used in more typical analyses; see Methods). Vector field learning was successful (**Figure S9C**). Fascinatingly, GBM cells were at the top of the combined cellular landscape (**Figure 7L**, **Figure S9D–E,** and **Supplementary Animation 3**). This finding suggests that overall the GBM cell state is more undifferentiated than normal stem cells within the developing brain. Most GBM cell types remained closely clustered at the top of this landscape with relatively similar Hodge potential (**Figure S9E-F**). These data suggest that GBM cell states are closely related and transitions between such cell states require less activation energy than transitions between normal developing cell states (**Figure 7M**). This indicates that GBM cells need to overcome lower energy barriers than normal brain cells to transition between cell states.

In support of this finding, we readily identified putative intermediate cell states via PhenoCycler (**Figure 7N** and **Figure S9G – I**), as defined by cells which are positive for both Vimentin (a MES-like marker) and OLIG2 (an OPC-like marker). Furthermore, we noted that the intermediate cells were often identified around OPC-like cells, suggesting a gradient of cell states at the interface between the two niches. By co-staining with Ki67, we verified that OPC-like and Intermediate cells are the predominant proliferative population in GBM and that cells in the core of the tumor are more proliferative than those at the periphery (**Figure S9J**).

If GBM cells readily transition between states, one would expect chromatin states at key genomic loci (those genes which control cell state and transition thereof) to be quite permissive to gene expression. Thus, we hypothesized that GBM cells exhibit high cell plasticity by keeping chromatin open at such genomic loci. To answer this question, we leveraged our single cell multiome dataset in which both RNA and ATAC data is generated for the same cell. After ordering the cells via RNA velocity, we divided them into four stages of pseudotime. As shown in **Figure S9K**, whereas gene expression for the key stem cell marker *NES* drastically decreased as cells progressed through pseudotime, chromatin remained accessible across the gene with only minor fluctuations, likely at distal regulatory elements.

Taken together, these data support a paradigm of GBM cell state plasticity, whereby the two major cell types of GBM can transition between one another at the spatial border of their respective niches. Furthermore, though other studies have suggested a high degree of cell plasticity in GBM, our data uniquely identifies a potential mechanism for this plasticity. We show that key transcription factors which specify GBM cell state (e.g. *FOS* and *ASCL1*) remain present and bound to DNA across GBM cell types, and that chromatin at key genomic loci (e.g. *FOS*, *NES*, *BCAN*, *BMP7, ANGPTL4*) remains open, facilitating rapid activation of transcription and cell state transition.

## DISCUSSION

Integrating spatial ATAC, spatial multiome, snATAC, and scRNA datasets generated from surgical specimens revealed the existence of two spatially segregated niches in GBM, which we named OPC-like and MES-like because (i) their molecular profiles were reminiscent of previously described OPC– and MES-like cell types and (ii) to avoid the introduction of unnecessary new nomenclature. OPC-like cells are characterized by accessible chromatin at neurodevelopmental genes and by programs associated with cycling cells. Chromatin programs of MES-like cells are dominated by the footprint of AP-1 TFs.

Our analyses revealed that OPC-like niches constellate and are enveloped by the larger MES-like niches. This spatial organization suggests crosstalk occurs between the two cell types, which we investigated further. We distinguished long– and short-range signals: those that operate within the niche, reinforcing its cell state (short-range signaling) and those that signal to the opposite niche, reinforcing that identity (long-range signaling). An example of long-range signaling is BMP-mediated signaling, which originates in the OPC-like niche and is directed to the MES-like niche. Based on our own data and related literature^58^, BMP-mediated signaling reinforces the MES-like state. An example of short-range signaling is PDGFB which is contained within the OPC-like niche, where it contributes to the maintenance of the OPC-like state. In summary, the spatial organization of niches therefore provides mechanisms to maintain self-states (short-range signaling) and to reinforce the neighboring state (long-range signaling).

Our two-niche model suggests OPC-like and MES-like cells are in a relationship of dynamic equilibrium, where MES-like cells can transition to the OPC-like state through an intermediate state. Our results are consistent with RNA velocity-based analyses that suggested MES to proneural state transitions^8^. Similarly, functional assays with cell state reporters show that the proneural/OPC-like state is stable – a ground state – while the MES-like state is metastable and easily able to transition to the OPC-like state^59^.

Our results are independently supported by orthogonal approaches, including scRNA-seq, PhenoCycler, and functional experiments with patient-derived models. What are the mechanisms that control this equilibrium? One may be geographic distance. Given that OPC-like niches tend to be small and very localized and are completely surrounded by much larger MES-like niches, a MES-like cell may need to be very close to an OPC-like niche to feel the effect of PDGFB produced by this inner niche. If the MES-like cell is far enough, it will instead be under the influence of the long-range signaling from the OPC-like niche and the short-range signals from the MES-like niche, both of which reinforce the MES-like state. Therefore, only a small fraction of MES-like cells located at the transition zone can transition to the OPC-like state. The two-niche model built through integration of highly complementary approaches simply explains the cellular organization of GBM and is consistent with a dynamic equilibrium between cell states that accommodates bidirectional cell state transitions.

This model of mutually-reinforcing (and therefore mutually beneficial) extracellular signaling-based GBM cellular niches coupled with a high degree of GBM cell type plasticity makes good sense of several previously published observations including (i) the transition of GBM cell states upon manipulation of cell culture media^44,59^ or implantation into the mouse brain^9^ (ii) the advantage of transplanting different GBM cell types together to produce the most rapid growth^60^, and (iii) cell state transitions under the selective pressure of cancer-directed treatments in patients^17^.

The current view of GBM tumor growth is based on the idea that the core of the tumor is composed of MES-like cells that are highly interconnected with each other, whereas the tumor periphery is populated by proneural/OPC-like cells that are unconnected, infiltrate the brain parenchyma, and proliferate^38,39^. Our findings help refine this view. We think that the infiltrative proneural/OPC-like cells act as guides for the expansion of the MES-like-dominated tumor mass. We speculate that GBM grows through two forces: (i) local cell proliferation in OPC-like niches in the tumor core, and (ii) centrifugal expansion guided by proneural/OPC-like cells invading the brain. This dual model of expansion does not require conversion of the OPC-like cells at the margins into MES-like cells, agrees with the increased proliferation propensity of OPC-like cells that has been reported by multiple groups and explains the metastable identity of MES-like cell states by invoking transition into OPC-like states.

The spatial niches we identified reflect the coexistence of true epigenetic/chromatin states and do not seem to reflect segregation of genetic subclones. In fact, spatial ATAC data gave us a platform to study the topographical distribution of genetic subclones characterized by focal amplifications of key GBM oncogenes, such as *EGFR*, *MDM2* and *CDK4*. We found that genetic subclones usually occupy specific regions of the tumor, but at the same time they are dispersed in cartesian space across niches.

In conclusion, here we provide support for the existence of two spatially segregated niches in GBM that are defined by chromatin accessibility and transcriptional profiles. Our two-niche model explains cell state transitions of GBM cells which possess a high degree of cellular plasticity.

## ACKNOWLEDGMENTS

This work was supported by Canadian Institutes of Health Research (CIHR) project grants (PJT-156278 and PJT-173475) and a Canada Research Chair to MG; and by National Institutes of Health (NIH) grants (R01CA245313, U54CA274509, U54CA268083) to RF.

## AUTHOR CONTRIBUTIONS

MG, RF and AN designed the study. SK, STY, SB, GL, DW, ES, XL, MZ, DZ, GS, AE, MY, HMC, KE executed experimental activities. CA, JM, JK and JAC provided samples and performed pathology annotations and quality control. AS and JZ supervised trainees and contributed to experimental design. SK, STY, SB, GL, AN, RF and MG contributed to writing this paper.

## MATERIALS AND METHODS

### Sample Preparation

Patient samples were obtained from primary GBM IDH-WT patients that had tumor resection performed in Foothills Medical Centre (Calgary, AB, Canada) or Yale New Haven Hospital from Feb 2019 to Jan 2021. All samples were collected and used for research with appropriate informed consent and with approval by the Health Research Ethics Board of Alberta (study ID HREBA.CC-16-0823) and Yale University Institutional Review Board (study ID 9406007680). All patients provided a signed informed consent. After resection, tissue pieces were cut into ∼5-10 mm^3^ and dropped in 1 mL of cryopreservation media [12.5% BSA and 10% DMSO in 600 µL DMEM (ThermoFisher Scientific, Catalog # 11995073) and 400 µL NS complete media (NeuroCult™ NS-A Proliferation Kit (Human) (StemCell Technologies, Catalog # 05751) supplemented with 10 ng/µL basic fibroblast growth factor (StemCell Technologies, Catalog # 78003.2), 2 ng/µL Heparin (StemCell Technologies, Catalog # 07980) and 20 ng/µL epidermal growth factor (StemCell Technologies, Catalog # 78006))]. Samples were subsequently slowly frozen (–1°C/minute) to –80°C in a CoolCell in and then transferred to liquid nitrogen for long-term storage.

Selected tissue pieces were thawed on ice, washed in ice-cold DMEM (Wisent Bioproducts, Catalog # 319-005-CL) for 5 minutes and subsequently washed in PBS before embedding in Tissue-Tek® O.C.T. Compound (Sakura, 4583) and snap-frozen in isopentane that was pre-chilled in liquid nitrogen. OCT embedded samples were subsequently cryosectioned at 10 µm at –20°C, H&E stained to evaluate the tissue intactness, tissue quality and presence of tumor in the tissue blocks and then stored at –80°C until further use. For H&E staining, slides are baked for 1 minute @37°C, followed by 30 minutes of fixation at –20°C in prechilled methanol. Fixed sections are subsequently rehydrated in ethanol series, stained for 4 minutes in hematoxilin, 10 seconds in 0.2% ammonia water and 10 seconds in alcoholic eosin (H&E staining).

### Transfer of pathologist annotations to Spatial Epigenomics

All code to reproduce the registration of the H&E on top of the BSA image are published in the following github repository. It also includes everything necessary to extract the annotations and to use them to classify the spatial epigenomic arrays.

### Spatial ATAC-seq

#### Sectioning & staining

OCT-embedded frozen tissue was equilibrated for ∼30 minutes in a prechilled cryostat at –20°C. The sections for spatial analysis were cut at 10µm thickness and placed in the middle of a SuperFrost plus slide (VWR, Catalog # 48311-703), and immediately after sectioning, the slides were placed at –80°C until further use. To determine the ROI for each sample, an H&E staining was performed on an adjacent section. For H&E staining, slides are baked for 1 minute @37°C, followed by 30 minutes of fixation at –20°C in prechilled methanol. Fixed sections are subsequently rehydrated in ethanol series, stained for 4 minutes in hematoxylin, 10 seconds in 0.2% ammonia water and 10 seconds in alcoholic eosin.

#### Section annotation and ROI selection

H&E sections were reviewed by a board-certified neuropathologist (AN) and assessed for cellularity, viability, and selection of suitable regions of interest for downstream processing. H&E sections were also scanned using an EVOS FL Auto 2 microscope (Thermo Fisher Scientfic) using automated tiling. Scanned sections were annotated in ImageJ by a staff neuropathologist (AN) and trainee neuropathologist (ELS), and annotations were set up as regions of interest in ImageJ.

#### Sequencing

Spatial ATAC-seq was performed according to the AtlasXomics protocol with a minor adaptation: before fixation, sections are baked for 5 minutes at 37°C instead of thawing for 10 minutes at room temperature. In short, tissues are fixed in 0.2% formaldehyde, and fixation is quenched with 1.25 M glycine, followed by a permeabilization step for 15 minutes at room temperature with 0.01% NP40. Next, in situ tagmentation was performed for 30 min at 37°C. Subsequently, ligation of spatial barcoded oligonucleotides was done using microfluidic chips with 50 channels of 25 µm thickness. First, microfluidics chip A was applied over the region of interest of the tissue, and ligation mix with channel-specific barcode A oligonucleotides for ligation was flown in each channel and incubated for 30 min at 37°C. Next, an identical second in situ ligation step of barcode B oligonucleotides was performed using a microfluidics Chip B, in which the channels flow perpendicular to the chip A channels over the region of interest. This results in a grid of 2500 unique barcode A-Barcode B combinations in tixels of 25 µm x 25 µm resolution. After ligation, tissue was lysed overnight at 40°C, and the lysate was cleaned using the DNA Clean & Concentrator-5 kit (Zymo research, Catalog # D4013). ATAC-seq libraries were prepared by amplifying the lysate using a non-indexed i5 and indexed i7 primer, and the optimal number of PCR cycles was determined using qPCR. Finally, libraries were sequenced paired-end 150 cycles on a Novaseq 6000 device (Illumina), aimed for ∼200 million read pairs per sample.

#### Analysis ATAC-seq

The raw sequencing data were demultiplexed using bcl2fastq v2.20. Fragment files were generated using the preprocessing software from (fastq2frags, AtlasXomics). In short, ligation linkers are filtered from Read2 using bbduk (bbmap), and alignment to GRCh38 genome is performed using Chromap.

Pathological annotations were made based by two independent neuropathologists on an H&E staining performed on a consecutive section for each sample. Annotations were exported using a custom IJM macro into a labeled image where each unique integer refers to a specific annotation. The labeled objects were then turned into a geojson file representing the polygons of the annotations, such that the coordinates can be transposed along with the H&E Image.

For each sample, a tailored variation of image processing operations is applied to match the H&E to the BSA image captured during the spatial barcoding. These include rotating, flipping, or cropping the image, with specific parameters that vary across samples (Supp. table X), using an in-house codebase. Afterwards, the H&E image was registered onto the BSA image. If the two images were sufficiently similar, automatic linear registration was used. In cases where this approach was inadequate, corresponding marker points were manually flagged, and the optimal transformation was computed based on these points. In both cases, warping the H&E into the coordinate space of the BSA was done using an affine transformation and a mutual information loss. If necessary, a second automatic registration step was performed allowing for nonlinear transformations to correct for tissue folding. All operations were mirrored on the previously mentioned polygon representations of the annotations.

Next, the spatial barcodes were projected on the BSA image using the AtlasXbrowser software (AtlasXomics) which uses the fluorescently labeled outer tixels as reference to project a 2D grid onto the image. This grid was used to define on– and off-tissue barcodes, and eventually, this creates a tissue position list with xy-coordinates per barcode. The AtlasXBrowser software was adapted such that the bounding box of its cropping functionality were saved as metadata along with the rotation, such that the same steps can be applied to the annotations, after which the transformed annotations were used to annotate the tixels, assigning a class to each barcode in the tissue position list.

Downstream analysis (QC, integration, spatial projection, etc.) was performed using ArchR^61^ and Seurat/Signac^62^.

#### TF motif enrichment

To assess TF motif enrichment in spatial ATAC data, we used ChromVAR^63^ as part of the Signac RunChromVAR package using the following parameters: (object = spatial_atac.object, genome = BSgenome.Hspaiens.UCSC.hg38). Enrichment scores were then calculated using the ChromVAR matrix as input and Seurat’s FindAllMarkers function with default parameters.

#### Gene set enrichment analysis

Differential gene activity results were sorted by log_2_FC in descending order. The resultant table was output to a .rnk file and used as input against either standard gene sets from the GSEA database or a .gmx file of the top 50 genes described to be enriched in Neftel cell types^9^.

#### Copy number computation and cellularity assessment

To assess copy number and cellularity in our samples, we extended our previously developed Copy-scAT R package to function for spatial samples^64^. We retained the raw signal pileup and normalization steps from Copy-scAT, but instead of collapsing adjacent intervals together, we applied CpG content-based normalization on each interval individually. We then generated a pseudobulk profile across the entire genome and used combined mean and variance changepoint analysis in the changepoint package with penalty = “MBIC” and method = “PELT” and a minimum segment length of 2 to identify individual segments, followed by estimation of copy numbers using a log-ratio approach. This approach enabled identification of large chromosome-level events and smaller local amplification regions. For assessment of cellularity, we used the imputed copy number of chromosome 7q as a proxy of tumour content. To calculate cellularity, we modelled the chromosome 7q estimated number in each sample using a two-component gamma distribution in the mixtools package with a simulated set of expected normal control spots. An adjustment factor was applied to account for different signal dispersion levels in samples with higher or lower signal to noise ratios. This model was used to generate an estimate of the expected chromosome 7q number in 100% tumour areas, which was then used to estimate the percent tumour content in each spot.

#### Weighted gene-network correlation analysis of spatial ATAC data (WGCNA)

Normalized gene score matrix data from ArchR was used to perform weighted gene-network correlation analysis (WGCNA) in the following manner. In a manner similar to hdWGCNA and scWGCNA [citations], the analysis was performed on pseudospots to improve per-cell sequencing depth. First, neighourhood clustering was performed using the Seurat method in ArchR with resolution = 5, with the goal of clustering the spots into approximately 25-35 ‘pseudospots’ per the high-quality thresholded spots on each slide. Samples with greater than 40 clusters were reclustered with iterative reduction of resolution by 0.75 x resolution until they had less than 40 clusters. Normalized pseudospots from all samples were combined, and WGCNA was performed with the following settings: corType = “pearson”, power = 12, networkType = “signed”, deepSplit=4, detectCutHeight = 0.995, pamRespectsDendro = F, minModuleSize = 10, maxBlockSize = 4000, TOMDenom = “min”, reassignThreshold = 1e-12, minKMEtoStay = 0, minCoreKME = 0.50, minCoreKMESize = 10, mergeCutHeight = 0.05, pamStage = T. Subsequently, all clusters with distance less than 0.15 were merged in a second merging step using the mergeCloseModules function in WGCNA. Consensus modules were generated by assigning each gene to the module with which it had the highest correlation score and restricting modules to genes with a minimum correlation threshold of 0.6. Module annotation was performed using a combination of gene enrichment analysis, correlation to annotated single-nucleus ATAC datasets generated in our laboratory, correlation to other published signatures^9^, as well as manual curation.

#### Spatially variable gene analysis and Moran I calculation

Analysis of spatially variable genes and assessment of autocorrelation via global and local Moran I was performed with Voyager^65^ in R 4.4.1. Spatial ATAC-seq gene score matrices were combined with spatial coordinates and used to generate a Voyager object. Neighbourhood graphs were constructed using the tri2nb function from the spdep package. Highly variable genes were generated with scran, and Moran I was computed for the top 1000 highly variable genes. Hierarchical clustering of Moran I score distances using the ward.D method was used to calculate distinct gene autocorrelation clusters. For computation of autocorrelation for module signatures or amplification scores, the same procedure was followed except using modules or amplification signatures as a Voyager assay instead of gene scores.

#### Neighbourhood domain analysis

For calculation of spatial domain size and territory, a custom function was built using the Voyager object as a scaffold. For each sample, the targeted domain score was thresholded using a set threshold (i >= 8 for amplifications, or i >= 6 for cortical signatures), and individual domains were generated by merging positive cells across contiguous neighbours using a distance-based neighbourhood graph (dnearneigh from spdep). Results were combined on a per-sample level and plotted in R.

### Spatial co-profiling

Spatial coprofiling involved in situ ATAC-seq and in situ gene expression (GEX) profiling simultaneously, followed by ligation-based barcoding using the AtlasXomics microfluidic chips and barcodes. For this, the AtlasXomics protocol for ATAC-seq was adapted based on a coprofiling protocol^22^. In short, frozen sections are baked for 5 minutes at 37°C, tissues are fixed in 0.2% formaldehyde, and fixation is quenched with 1.25 M glycine, followed by a permeabilization step for 15 minutes at room temperature with 0.01% NP40. Next, in situ tagmentation was performed for 30 minutes at 37°C, followed by in situ reverse transcription with biotinylated primers. Subsequently, ligation of spatial barcoded oligonucleotides was done using microfluidic chips with 50 channels of 25 µm thickness. First, microfluidics chip A was applied over the region of interest of the tissue, and ligation mix with channel-specific barcode A oligonucleotides for ligation was flown in each channel and incubated for 30 min at 37°C. Next, an identical second in situ ligation step of barcode B oligonucleotides was performed using a microfluidics Chip B, in which the channels flow perpendicular to the chip A channels over the region of interest. This results in a grid of 2500 unique barcode A-Barcode B combinations in tixels of 25 µm x 25 µm resolution.

After ligation, tissue was lysed overnight at 40°C, and the lysate was cleaned using the DNA Clean & Concentrator-5 kit (Zymo research, Catalog # D4013). Biotinylated RNA-cDNA hybrids were separated from transposed DNA fragments for ATAC-seq by Streptavidin-coated magnetic beads. The supernatant was cleaned using DNA Clean & Concentrator-5 kit (Zymo research, Catalog # D4013) and used for ATAC-seq library preparation by amplifying the lysate using a non-indexed i5 and indexed i7 primer, and the optimal number of PCR cycles was determined using qPCR. The bound fraction was used for on-bead template-switching and cDNA pre-amplification, followed by cDNA amplification of the supernatant. 1ng of amplified cDNA was used for library preparation using Nextera XT library prep kit (Illumina) with custom non-indexed i5 and indexed i7 primers. Quality of cDNA, GEX libraries and ATAC-seq libraries was checked on a TapeStation D5000 screentape system (Agilent, Catalog # 5067-5588 & 5067-5589) and finally, libraries (ATAC-seq and GEX) were sequenced paired-end 150 cycles on a Novaseq 6000 device (Illumina), aimed for ∼200 million read pairs per sample.

#### Analysis co-profiling

The raw sequencing data were demultiplexed using bcl2fastq v2.20. ATAC-seq data were analysed as described above. GEX data were first preprocessed with a variant of the AtlasXomics pipeline.

Both data layers (ATAC-seq and GEX) were processed with a variant of the AtlasXomics preprocessing pipeline, where ligation linkers are filtered from Read2 using bbduk (bbmap), followed by read 2 separation into Read 2 and Read 3 with a custom script (from AtlasXomics, bc_process.py) and alignment of the reads was subsequently performed to the hg38 reference genome for the Multiome pipeline (GRCh38 – 2020-A, 10x Genomics) and quantified using cellranger-arc count function with default parameters (Cell Ranger ARC, version 2.0.2, 10x Genomics), yet with adapted barcode file.

#### NicheCompass Analysis

RNA and ATAC raw data were loaded into Seurat/Signac and reformatted to .h5ad files for input into NicheCompass. Gene programs and the subsequent analysis pipeline were run with default parameters as described (https://nichecompass.readthedocs.io/en/latest/)^37^.

### Single nucleus ATAC

#### Data analysis

Fastq files from snATAC experiments (10x Genomics Chromium Next GEM Single Cell ATAC Kit) from three patients were obtained from a previous paper^64^. Tumor chunks from these patients were also used for spatial ATAC profiling. scATAC-seq reads were aligned to the hg38 reference genome for the Multiome pipeline (GRCh38 – 2020-A, 10x Genomics) and quantified using cellranger-atac count function with default parameters (Cell Ranger ATAC, version 2.1.0, 10x Genomics). Downstream analysis was performed using Seurat/Signac^62^.

Quality control parameters were calculated using the NucleosomeSignal and TSSEnrichment functions of Signac. Quality control filters were as follows: gbm.atac, subset = nCount_ATAC > 10000 & TSS.enrichment > 3 & blacklist_ratio < 0.01 & nucleosome_signal < 2 & pct_reads_in_peaks > 25. Samples were batch corrected using Harmony^66^.

For cluster assignment, the neighborhood graph was constructed using the FindNeighbors function with the following non-default parameters: reduction = ‘harmony’, dims = 2:30. Clusters were called using FindClusters with the following non-default parameters: algorithm = 3, resolution = 0.4. Gene activity scores were calculated with GeneActivity function using default parameters, and clusters were annotated based on review of markers from the FindAllMarkers function.

#### CellChat

Analysis of cell-cell communication was conducted using CellChat^42^. The gene activity matrix was used as input. Communication probabilities was calculated using computeCommunProb using the following parameters: type = “triMean.” The rest of the analysis pipeline was conducted using default parameters.

#### Gene set enrichment analysis

Gene set enrichment was performed using the GSEA graphical user interface application^67^. Differential gene activities from OPC-like and MES-like cells were filtered to only include those with an adjusted p-value of less than 0.05. The resultant table was then sorted by log2FC and output to a .rnk file. The run GSEAPreranked tool was used. In addition to the standard databases available through GSEA, enrichement was also run against a custom .gmx file which contained the top 50 genes upregulated in the Neftel, et al. cell types^9^.

#### TF Motif Enrichment

Differential peaks were identified between OPC-like and MES-like cells using the FindMarkers function with the peaks matrix as input using default parameters. Results were then filtered to only include statistically significant peaks (adjusted p vale < 0.05). Upregulated peaks (present in OPC-like) and downregulated peaks (present in MES-like) were split into two distinct tables. Motifs enrichment was then calculated using the FindMotifs function with the feature input being the coordinates of the differentially accessible peaks. TF footprints were visualized using the Footprint function with the following parameters: genome = BSgenome.Hsapiens.UCSC.hg38, in.peaks = TRUE.

#### Gene regulatory network

To ascertain the putative gene regulatory networks of TFs of interest, we first identified promoter and enhancer regions for targets of interest using the GeneHancer database^50^. Coordinates were output as a .bed file, and sequences from those coordinates were then recovered using the bed2fasta function of MEME Suite^68^. Motifs matrices for TFs of interest were retrieved from the JASPAR database^69^. Those sequences were then analyzed for motif enrichment using FIMO with the following parameters: –-bgfile –nrdb--–-thresh 1.0e-4. For ease visualization of these sites along ATAC coverage plots, the coordinates were expanded +200 bp on either side prior to plotting the region using the CoveragePlot function.

### Single-cell RNA velocity analysis

#### Data analysis

FASTQ files from the Couturier and colleagues study were accessed through the European Genome-Phenome Archive (Accession Number: EGAD00001006206). To generate the spliced/unspliced matrix for downstream velocity analysis, we used velocyto^70^ using the FASTQ files as input. A loom file with the count, spliced, unspliced, and ambiguous matrices as layers was generated using Seurat and SeuratDisk.

Cells were filtered according to the following quality control parameters: nFeature_RNA > 500 & nFeature_RNA < 6000 & nCount_RNA < 10000 & percent.mt < 10. The dataset was batch corrected using Harmony. To call clusters, we first constructed a neighborhood graph using the following non-default parameters: reduction = “harmony”, dims = 1:20. Clusters were called using FindClusters with the following non-default parameters: reduction = “harmony”, resolution = 0.2. Clusters were then annotated using expression of key markers identified using FindAllMarkers. Only malignant cells were included in downstream analysis. Malignant cell types were defined using CellCycleScoring and AddModuleScore with previously published marker genes as inputs^9^.

#### Dynamo

Velocity analysis was conducted using dynamo-release^51^. Pre-processing was conducted using the Preprocessor class of dynamo with recipe = “Seurat”. The harmony values for cells were used rather than principal components given the significant batch effect present in the dataset. The vector field was calculated using dyn.vf.VectorField with the following non-default parameters: M = 1000, pot_curl_div = True.

To calculate cell state transitions, individual cells in the dataset must be selected. Based on UMAP coordinates, cells at the extremes of their respective cluster within the vector field were selected for each major GBM cell type.

For three-dimensional visualization of ddhodge potential, the potential was first inverted by multiplying by –1. This produces a more intuitive visualization of cell differentiation (earlier cells higher and differentiated cells lower). The 3D plots were then generated using Plotly.

Gene regulatory networks are graphical representations of the Jacobian (first derivative of the vector field equation; see publication by Qiu and colleagues for details^51^). For calculation of the Jacobian, the dataset was randomly downsampled to 4000 cells given constraints of the computational resources available. The top 2000 most variable genes were used, and the Jacobian was calculated with dyn.vf.jacobian with default parameters.

#### Developing brain and GBM Integration

Single cell RNA sequencing data, including spliced/unspliced counts, were downloaded from the Github repository of Braun and colleagues^57^. Data were processed as described above for the GBM dataset. To more easily conduct analyses with computational resources that were available, the data were randomly downsampled to 50,000 cells.

Seurat objects for GBM and developing brain were merged. Unintegrated dimensionality reduction was performed with principal components analysis. Canonical correlation analysis integration was then performed using Seurat’s IntegrateLayers function with the following parameters: method = CCAIntegration, orig.reduction = “pca”, new.reduction = “integrated.cca”, dims = 1:50, dims.to.integrate = 50. We then joined count and data layers using JoinLayers. Nearest neighbor graph and UMAP dimensionality reduction was then performed on the integrated.cca vectors.

To perform vector field calculations on this dataset, we imported the CCA vectors for each cell into the dynamo Anndata object as the principal components (adata.obsm[“X_pca”] = cca_array). Downstream dynamo analysis was then performed as previously described.

### Single-cell multiome

#### Tissue dissociation

Single-cell suspensions were prepared as described previously^71^. In short, a cryopreserved tissue chunk of ∼5-15 mm^3^ was thawed on ice in pre-chilled Homogenization Buffer [5 mM CaCl_2_, 3 mM Mg(Ac)_2_, 10 mM Tris pH 7.8, 0.0167 mM PMSF, 0.167 mM DTT, 320 mM sucrose, 0.1 mM EDTA, 0.1% NP40, 1 U/µL protector RNAse inhibitor (Roche, Catalog # 3335399001)] and subsequently dissociated into single nuclei using a disposable pellet pestle. This homogenate is strained through a 35 μm cell strainer fluorescence-activated cell sorting tube (Thermo Fisher Scientific, 08-771-23) to remove debris, and then mixed in a 1:1 ratio with Iodixanol 50% buffer [4.25 mM CaCl_2_, 2.55 mM Mg(Ac)_2_, 8.5 mM Tris pH 7.8, 0.014 mM PMSF, 0.14 mM DTT, 50% Iodixanol Solution (OptiPrep, STEMCELL Technologies, Catalog # 07820), 1 U/µL protector RNAse inhibitor (Roche, Catalog # 3335399001)]. Subsequently, 600 µL of 35% and 29% Iodixanol solutions [5 mM CaCl_2_, 3 mM Mg(Ac)_2_, 10 mM Tris pH 7.8, 0.0167 mM PMSF, 0.167 mM DTT, 1 U/µL protector RNAse inhibitor, 35% or 29% Iodixanol Solution (OptiPrep, STEMCELL Technologies, Catalog # 07820)] were layered in a 2 mL microcentrifuge tube, followed by 400 µL of 25% Iodixanol solution containing the nuclei. The tube was spun for 40 minutes, 4500 RCF at 4°C in a swinging bucket centrifuge with lowest acceleration and deceleration settings to isolate the nuclei from any debris. The layer of nuclei was collected, and purity and concentration of nuclei was measured on a Countess II cell counter (Invitrogen) using trypan blue dye. 106 nuclei were aliquoted and washed using wash buffer [10 mM Tris-HCl pH 7.4, 10 mM NaCl, 3 mM MgCl_2_, 1% BSA, 0.1% Tween-20, 1 U/µL protector RNAse inhibitor (Roche, Catalog # 3335399001)], lysed using 0.1X lysis buffer [10 mM Tris-HCl pH 7.4, 10 mM NaCl, 3 mM MgCl_2_, 1% BSA, 0.01% Tween-20, 0.01% Igepal, 0.001% Digitonin, 1 mM DTT, 1 U/µL protector RNAse inhibitor (Roche, Catalog # 3335399001)] and washed again in 1 mL of wash buffer. Finally, nuclei were resuspended in 1X Nuclei buffer (10x Genomics, 2000207) supplemented with 1 mM DTT, in a concentration ranging from 2580 to 4650 nuclei/µL (aiming for 8000 nuclei in the subsequent 10x Genomics Chromium protocol) and counted using a Countess II cell counter (Invitrogen) with trypan blue dye to verify the concentration.

#### Library preparation

Resuspended nuclei are processed with the Chromium Next GEM Single Cell Multiome ATAC + Gene Expression Reagent Bundle (10x Genomics, Catalog # 1000285) according to manufacturers’ instructions. In short, 5µL of 2580-4650 nuclei/µL are used for transposition for ATAC-seq, followed by partitioning of single nuclei into Gel Bead-in-Emulsion (GEMs). In these GEMs, lysis, reverse transcription of poly-adenylated RNA and GEM-specific barcoding of the transposed DNA and full-length cDNA from the encapsulated cell happens. Next, GEMs are dissolved, and the pooled fraction is pre-amplified before splitting the sample for separate ATAC library preparation and cDNA amplification and subsequent GEX library preparation. Quality of cDNA, GEX libraries and ATAC libraries was checked on a TapeStation D5000 screentape system (Agilent, Catalog # 5067-5588 & 5067-5589) before sequencing on a Novaseq 6000 device (Illumina).

#### Initial data analysis

The raw sequencing data were demultiplexed using cellranger-atac mkfastq (Cell Ranger ARC, version 2.0.2, 10x Genomics). scATAC-seq and GEX reads were aligned to the hg38 reference genome for the Multiome pipeline (GRCh38 – 2020-A, 10x Genomics) and quantified using cellranger-arc count function with default parameters (Cell Ranger ARC, version 2.0.2, 10x Genomics).

#### RNA velocity

For RNA velocity analysis, the scRNA portion of multiome data was processed similar to the GBM dataset as described above. Calculated ddhodge potential was then divided into quartiles and each cell was assigned to one of those quartiles termed Early, Mid-Early, Mid-Late, and Late. This assignment was then added to the integrated Seurat object which contained ATAC and RNA data in different layers of the Seurat object. Coverage plots were generated with CoveragePlot.

### PhenoCycler

PhenoCycler-Fusion imaging was performed per the standard manufacturer’s protocol (Akoya Biosciences, https://www.akoyabio.com/wp-content/uploads/2021/01/CODEX-User-Manual.pdf). Briefly, fresh-frozen tissue sections (10 µm thickness) were mounted onto positively charged microscope slides. Slides were fixed in acetone for 10 minutes and 1.6% PFA for 10 minutes at room temperature followed by air drying and rehydration in the rehydration buffer. Slides were then incubated with a blocking solution provided by Akoya Biosciences for 30 minutes. Tissue sections were then incubated at room temperature for 3 hours with a panel of oligonucleotide-conjugated primary antibodies (Akoya Biosciences and in-house conjugated using Lightning-Link® Streptavidin Antibody Labeling Kit) targeting biomarkers of interest. Following incubation, slides were washed thoroughly and then post-fixed using 1.6% PFA and treated with ice-cold methanol before a final fixative step.

The PhenoCycler imaging process used iterative cycles of fluorescent dye-conjugated oligonucleotide reporters (Akoya Biosciences) to hybridize and visualize bound primary antibodies. Each imaging cycle included reporter dye hybridization, image acquisition, and subsequent dye removal to prepare slides for the next cycle. Images were acquired using an inverted fluorescence microscope with a 20X objective.

#### Data analysis

Image processing and analysis were conducted using Akoya Biosciences’ proprietary software and QuPath, which integrated data from multiple imaging cycles, enabling comprehensive spatial profiling and quantitative analyses. Cell segmentation was performed using InstanSeg^72^ for nucleus and/or cell segmentation in fluorescence images in Qupath v0.6^73^ using DAPI, VIM and OLIG2 as input channels. Next, DAPI+ segments were defined as cells, and tumor cells were identified as cells, excluding immune cells (CD45+), neurons (NEUN+), microglia (TMEM119^+^/IBA1^+^) and macrophages (CD68+). MES-like tumor cells were determined as VIM^+^; OPC-like tumor cells were determined as OLIG2^+^; and intermediate cells were determined as OLIG2^+^VIM^+^ cells. Next, cross-pair correlation function (cross-PCF) and its extension topographical correlation map (TCM)^74^ was used to identify spatial correlation between cell types.

### Evaluation of ligands on cell state

#### Cell Culture and Treatment

Patient-derived glioma stem cell lines GSC003 and GSC016 were cultured and seeded onto μ-Slide 8 Well Glass Bottom chambers (ibidi USA, 80827) pre-coated with poly-L-ornithine and laminin. For GSC003, 3 × 10⁴ cells were seeded per well; for GSC016, 2 × 10⁴ cells per well. Cells were allowed to adhere overnight prior to treatment with either 150 ng/mL rhBMP7 (R&D Systems, 354-BP-010/CF) or 500 ng/mL rhANGPTL4 (R&D Systems, 4487-AN-050) for 72 hours.

#### Immunocytochemistry

Following treatment, cells were fixed with 4% formaldehyde (Thermo Scientific, 28906) in Dulbecco’s Phosphate Buffered Saline (DPBS, pH 8.0) for 15 minutes at room temperature. Cells were then permeabilized and blocked using 5% Normal Goat Serum (Cell Signaling, 5425) in DPBS containing 0.3% Triton™ X-100 for one hour. Primary antibody staining was performed overnight at 4 °C using anti-OLIG2 (Cell Signaling, 65915) diluted 1:400 in DPBS supplemented with 1% Bovine Serum Albumin (BSA) and 0.3% Triton™ X-100 (pH 8.0). Following primary antibody incubation, cells were incubated with Alexa Fluor 647-conjugated goat anti-rabbit secondary antibody (Invitrogen, A21244), diluted 1:500 in the same buffer, for one hour at room temperature. Nuclear counterstaining was then performed using DAPI (Invitrogen, 62248), diluted to 2 μg/mL in DPBS (pH 8.0).

#### Imaging and Quantification

Images were acquired using a Nikon Ti2-E Spinning Disk Confocal Microscope equipped with Plan Apo λ 20X objective. For each well, three non-overlapping fields of view were imaged. For each field, 25 z-stack images were collected at 0.9 μm intervals. Maximum intensity projections were generated using ImageJ/Fiji, and brightness and contrast were adjusted uniformly across conditions for each cell line. Images were converted to 8-bit and processed using CellProfiler to quantify the percentage of OLIG2⁺ cells (children) within the total population of DAPI⁺ nuclei (parents). Cell counts from the three fields of view were summed to generate a single data point per well. Quantitative results were analyzed and visualized using R.

## FIGURE LEGENDS

**Figure S1.**
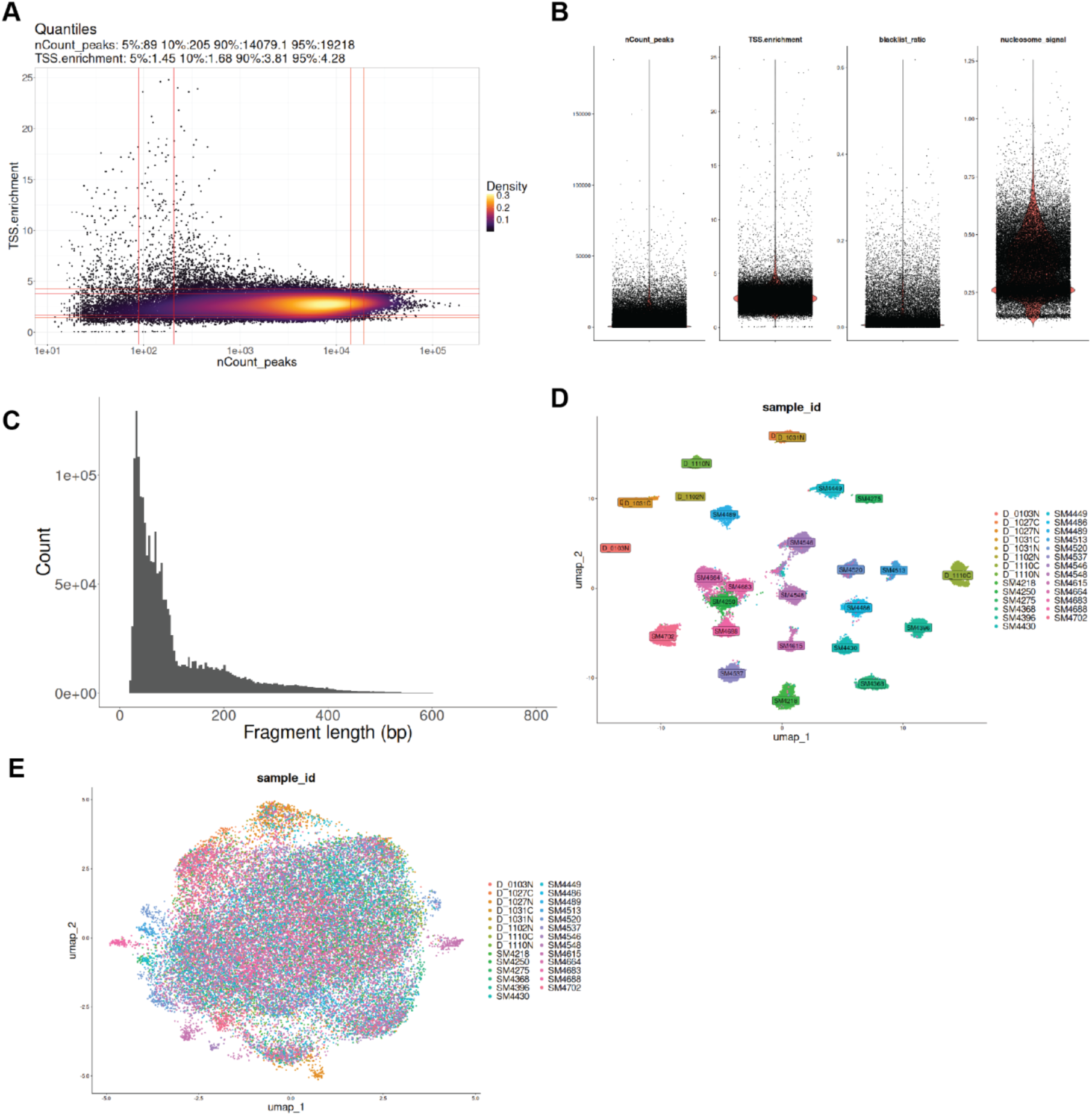
Quality control and overview of spatial ATAC data. Related to Figure 1. (A) Quality control data of spatial ATAC cohort showing quantiles of TSS enrichment and total number of peaks. (B) Violin plot of key quality control data. (C) Fragment histogram. (D) UMAP plotting of spatial ATAC pixels without batch correction. (E) UMAP plotting of spatial ATAC pixels with Harmony batch correction.

**Figure S2.**
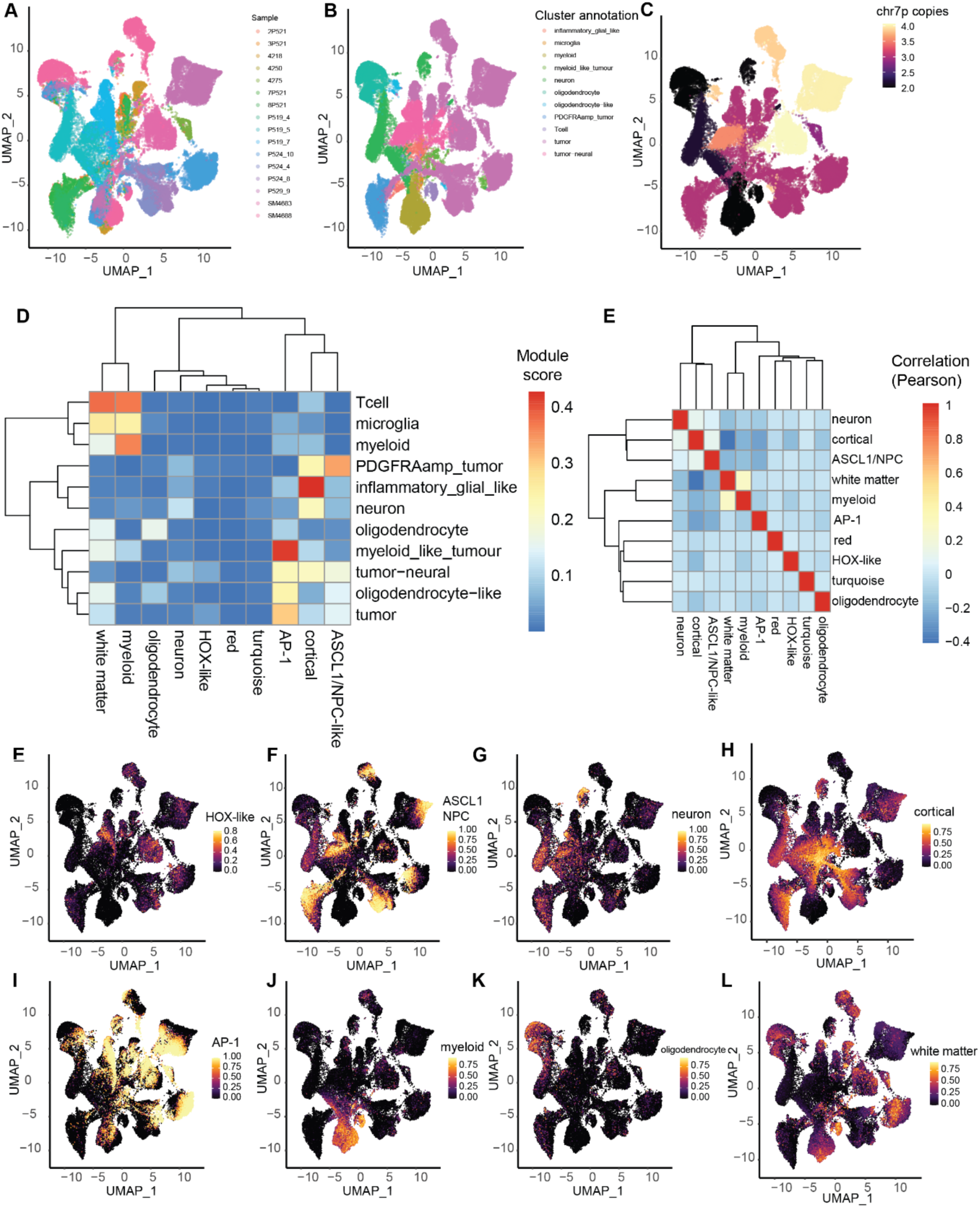
WCGNA module annotations. Related to Figure 2. (A) Overview of samples in combined dataset. (B) Annotation of cluster identities in a combined snATAC dataset. (C) Copy-scAT annotation of chromosome 7 copy number in different clusters. (D) Clustering of motif modules to different cell identities. (E) Pearson correlation of different module scores. (F – M) UMAP plot of snATAC datasets colored by indicated WCGNA module score.

**Figure S3.**
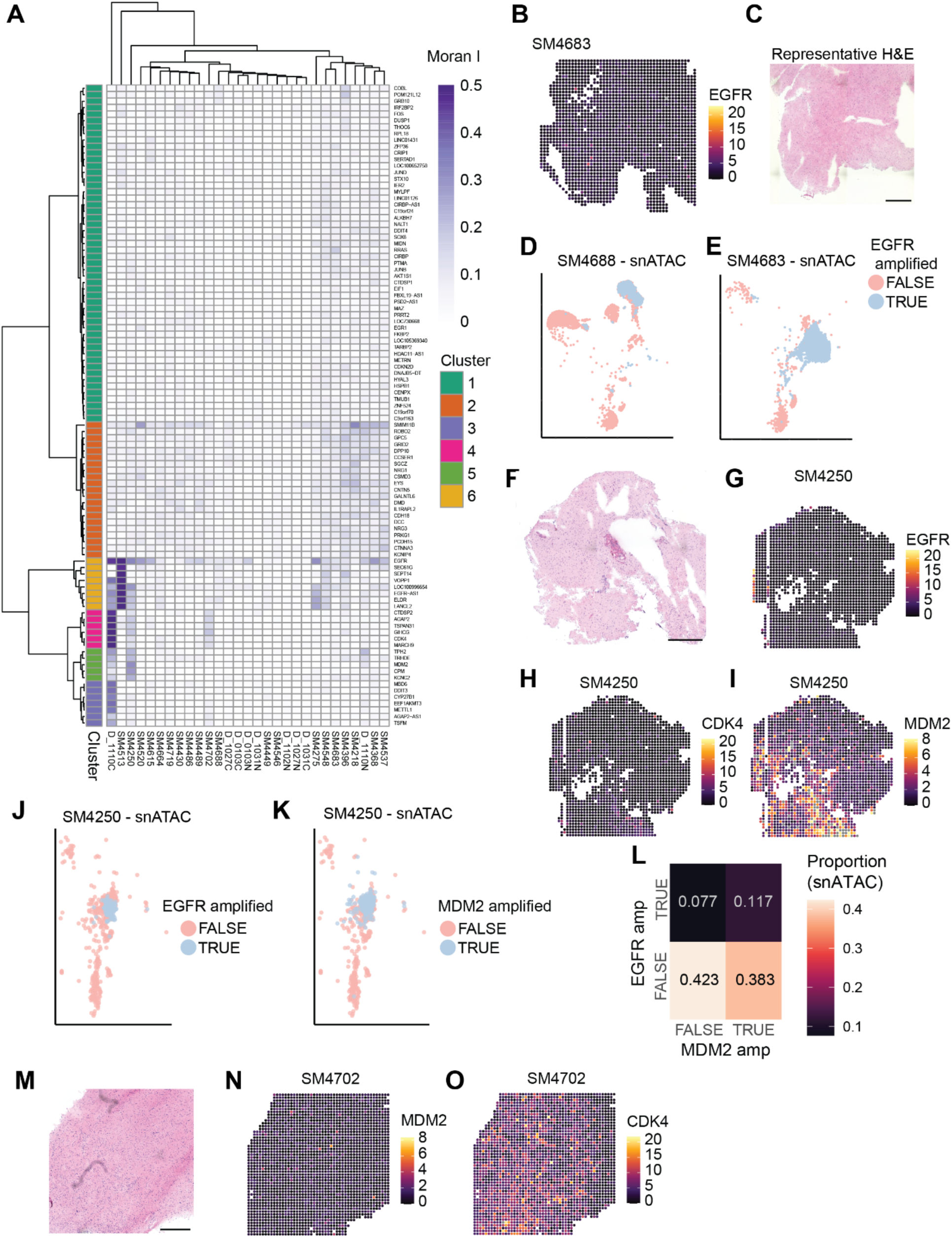
Subclonal genetic amplifications highlighted by spatial autocorrelation. Related to Figure 3. (A) Spatial autocorrelation plots for top 100 highly variable gene accessibilities. (B) Spatial plot of cluster 6 (EGFR amplification). (C) Representative H&E image for the sample in A. (D – E) UMAP plots of the snATAC data from the same sample colored by called EGFR amplification. (F – I) Spatial plots colored by the indicated gene amplification cluster along with representative H&E. (J – K) UMAP plot of snATAC data from the sample in F – I colored by called EGFR or MDM2 amplification. (L) Confusion matrix of EGFR and MDM2 amplification status based on snATAC data for sample SM4250. (M – O) Spatial plots colored by indicated gene amplification cluster and representative H&E.

**Figure S4.**
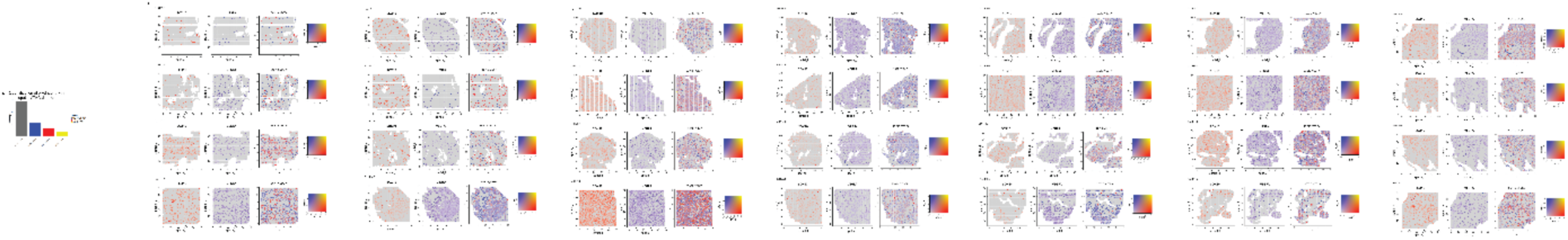
Spatial plots highlighting OPC-like and MES-like niches across the dataset. Related to Figure 4. (A) Quantification of pixel identity. (B) Spatial plots of all samples displaying gene activity for *SOX10* and *VEGFA*.

**Figure S5.**
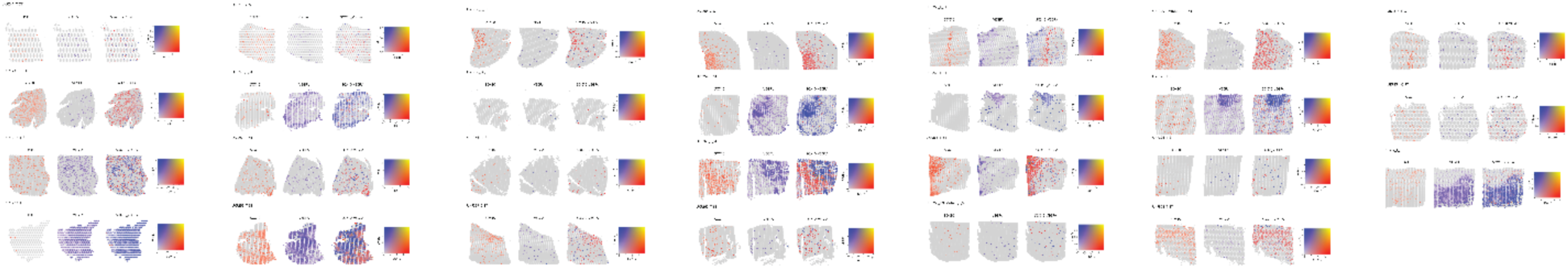
Spatial plots examining OPC-like and MES-like niches in independent Ravi, et al dataset. Related to Figure 4. Spatial plots from Ravi, et al showing transcriptomic quantification of *SOX10* and *VEGFA* genes.

**Figure S6.**
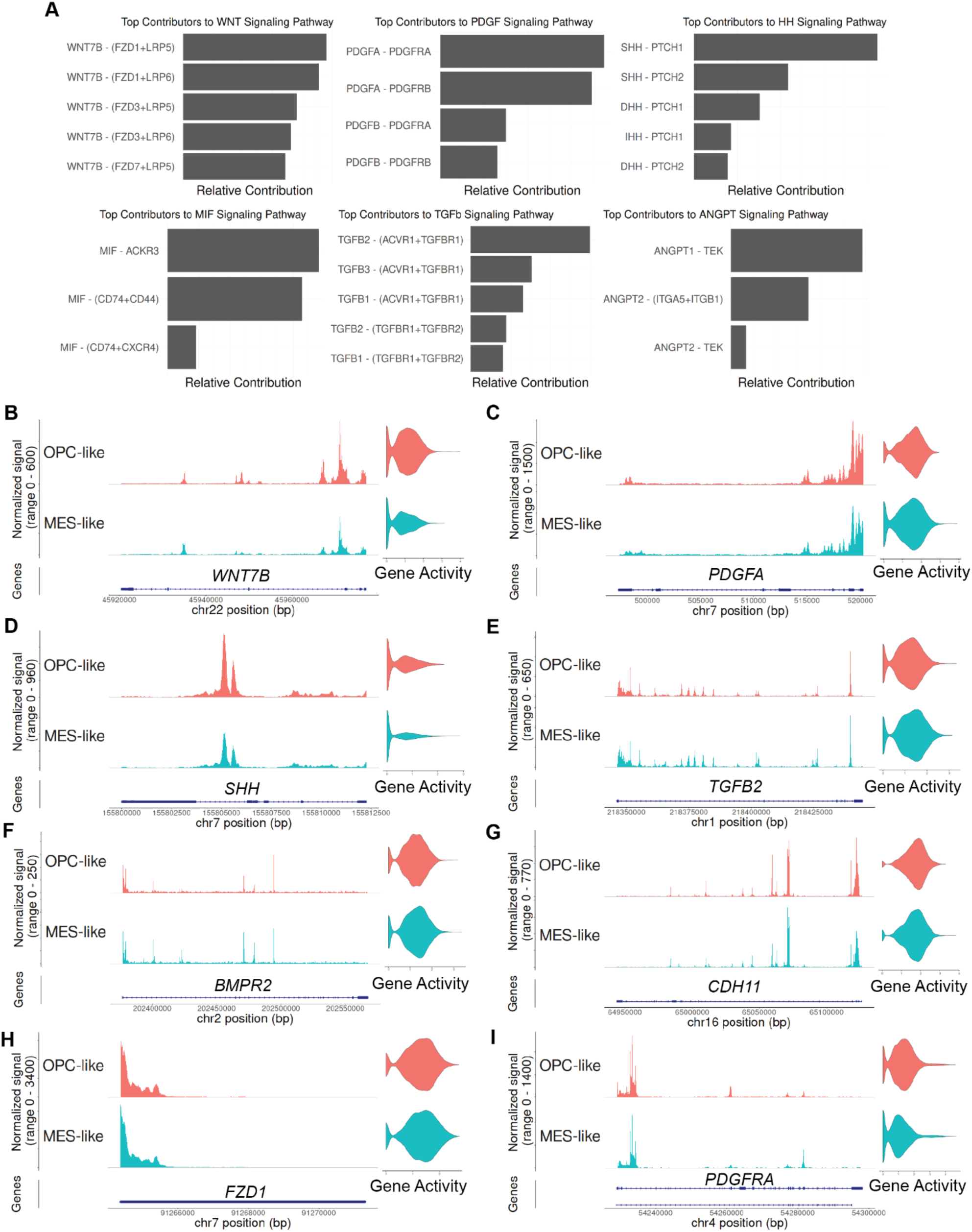
Overview of signaling programs active in GBM. Related to Figure 5. (A) Top contributing ligand-receptor pairs to the signaling pathways shown in Figure 5. (B – I) Coverage plots of top ligand and receptors for key pathways. Note the similarities in gene chromatin accessibility across cell types.

**Figure S7.**
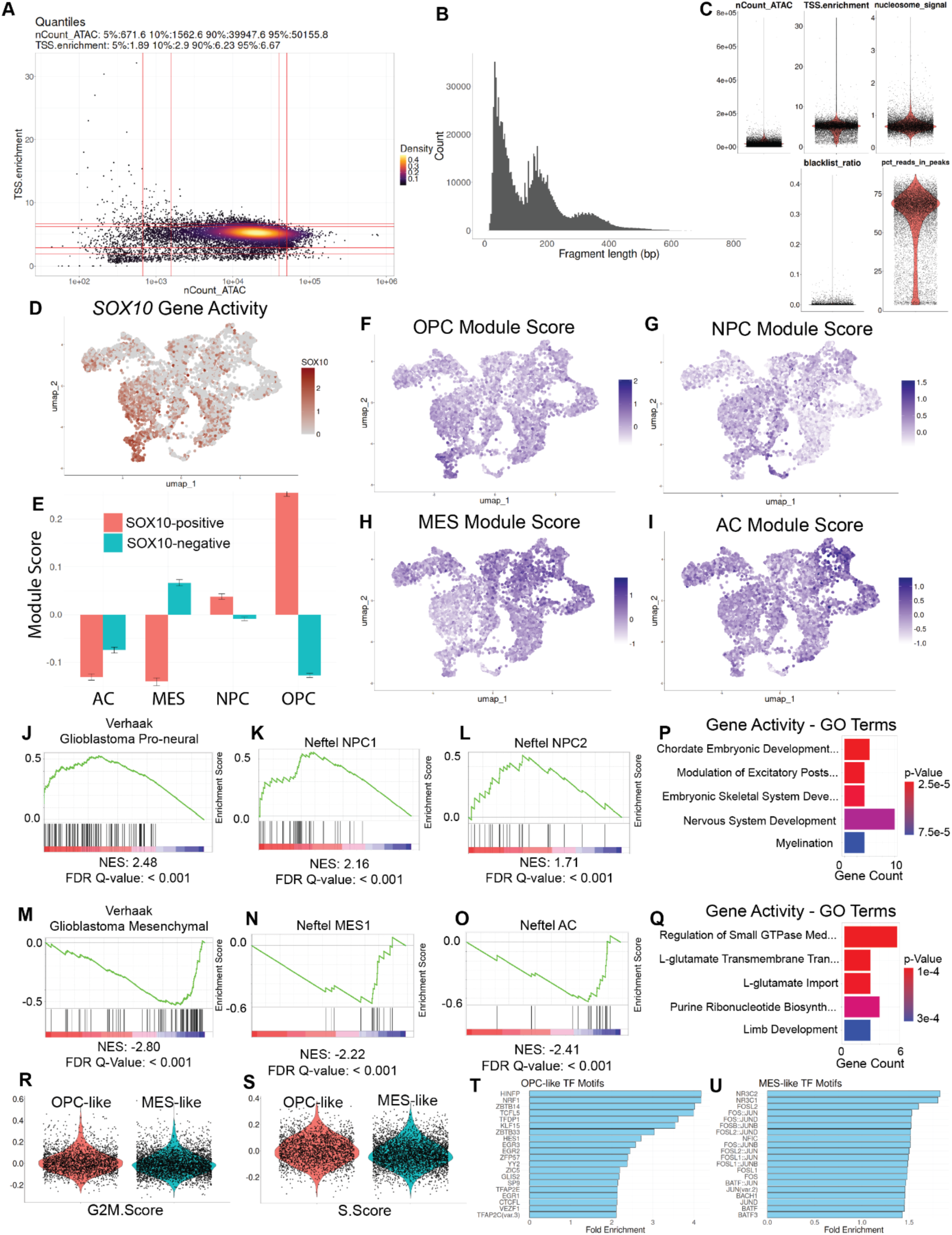
Quality control and annotation of single nucleus ATAC data. Related to Figure 6. (A) Number of peaks in single nucleus ATAC data plotted against TSS enrichment. Quantiles are marked. (B) Fragment histogram of snATAC data set. (C) Violin plots of key quality control metrics for snATAC data set. (D) UMAP plot of malignant cells colored by *SOX10* gene activity. (E) Quantification of module scores for key GBM cell types validating use of *SOX10* gene activity as a cell type marker. (F – I) UMAP plot of malignant cells colored by respective cell type module scores. (J – O) Gene set enrichment analysis of differential gene activities between SOX10-positive and SOX10-negative cells against known GBM tumor and cell types. (P – Q) Top five enriched GO terms for differential gene activities between OPC-like and MES-like cell types. (R – S) Violin plot of G2M and S module scores showing both higher mean score in OPC-like cells but with a distribution such that some MES-like cells are likely cyclin. (T – U) Bar graph demonstrating fold enrichment of TF motifs for OPC-like and MES-like cells. For all motifs shown, the p-value is less than 0.00001.

**Figure S8.**
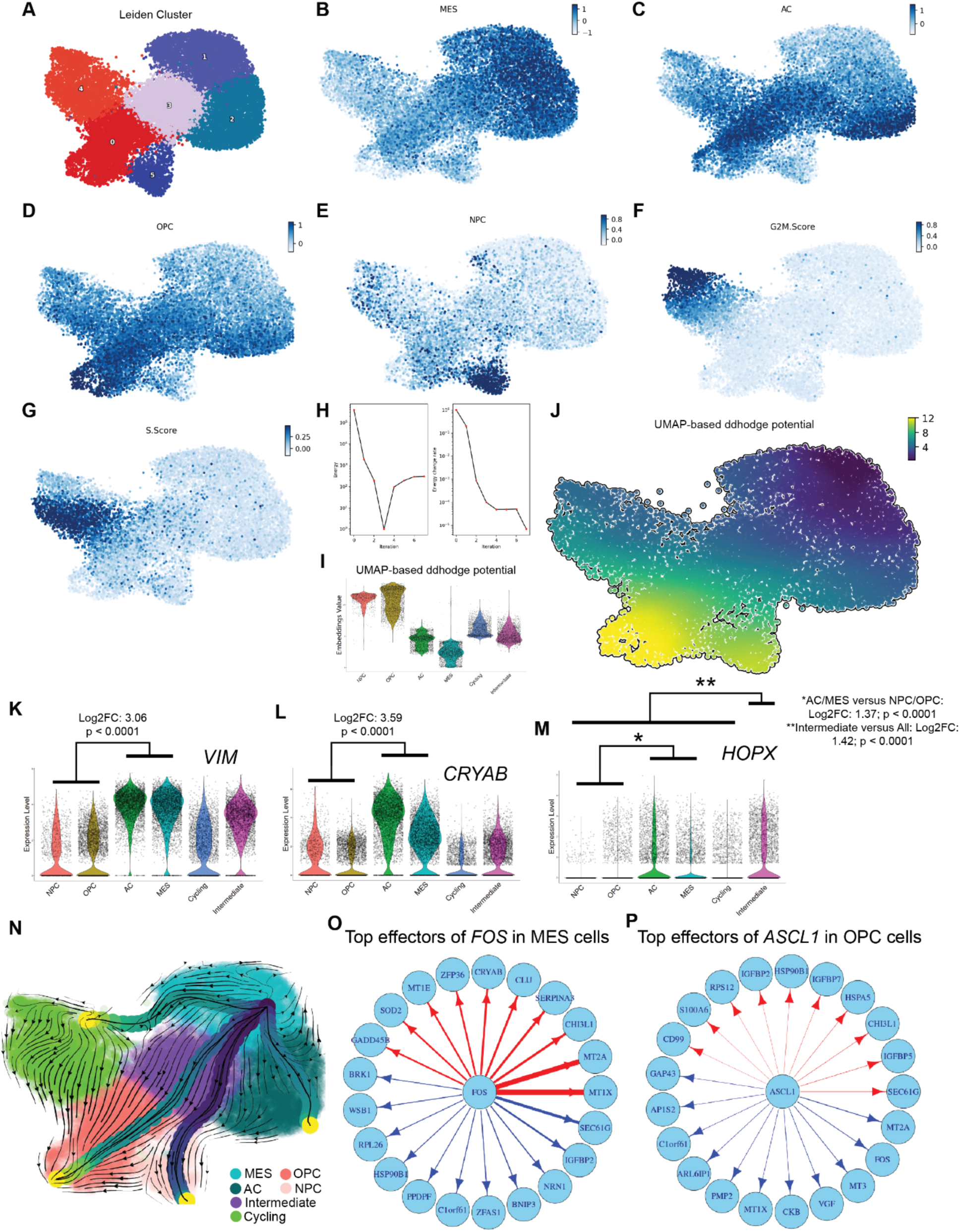
Analysis of RNA velocity data from Couturier, et al. Related to Figure 7. (A) UMAP plot of single-cell RNA sequencing data colored by Leiden clusters. (B – G) UMAP plot colored by module scores for indicated gene set. (H) Total energy and energy change across iterations as the vector field was learned. (I) Violin plot of ddhodge potential for major GBM cell types. (J) UMAP plot colored by ddhodge potential. (K – M) Violin plots of indicated gene expression with statistically significant differences of interest highlighted. (N) Least action paths for transition from MES cells to other cell types. (O – P) Network graph of putative gene regulatory networks for TFs of interest. Red indicates a positive (upregulation) relationship while blue indicates a negative (downregulation) relationship. The edge width indicates the strength of the calculated interaction.

**Figure S9.**
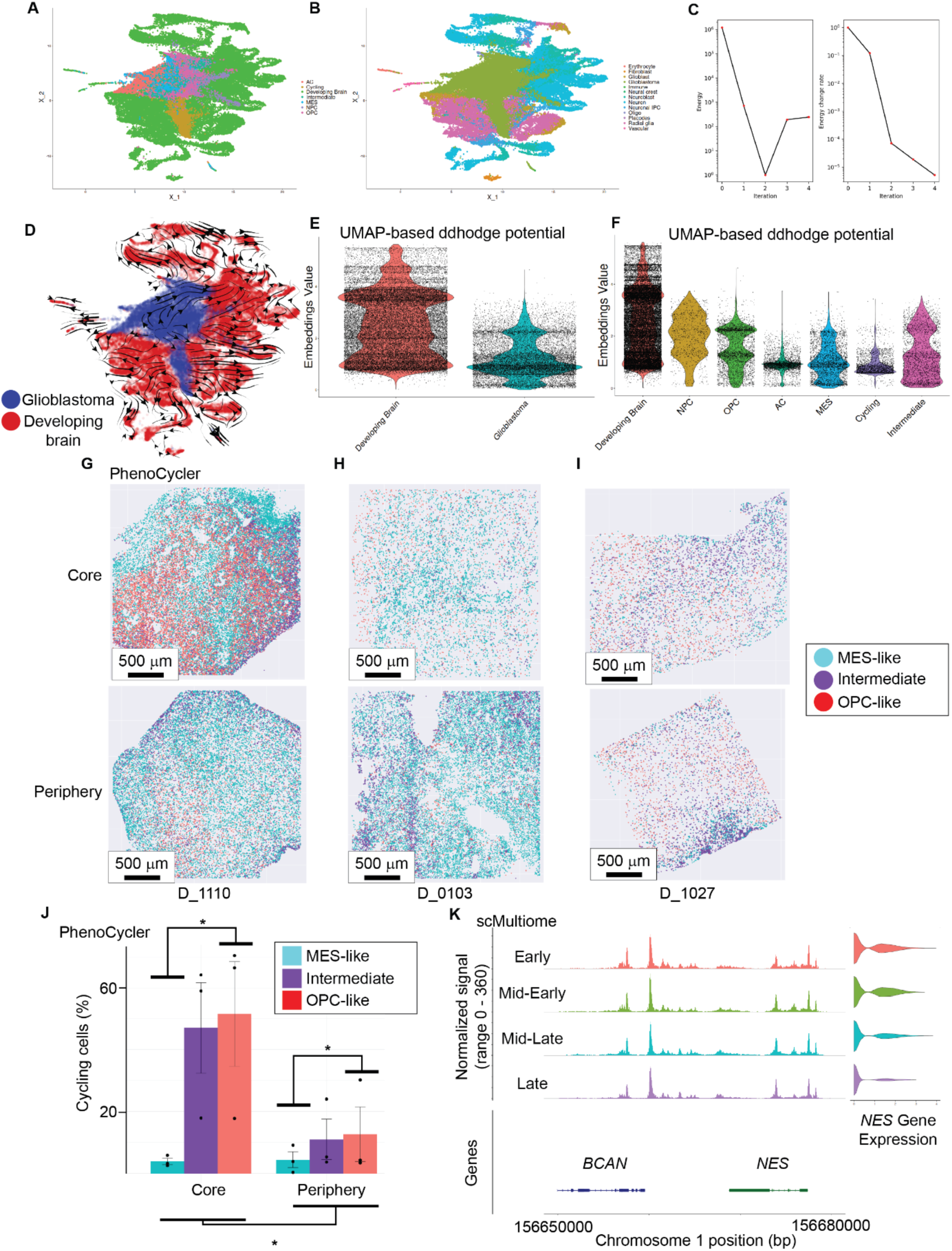
Analysis of combined GBM and developing brain with validation from PhenoCycler and multiome data. Related to Figure 7. (A – B) UMAP plot of integrated single cell RNA data from GBM and developing brain colored by indicated cell types. (C) Total energy and energy change across iterations as the vector field was learned. (D) Streamline plot of integrated GBM and developing brain. (E) Violin plot of ddhodge potential of normal developing brain versus GBM. (F) Violin plot of ddhodge potential of developing brain with GBM cells divided into the major GBM cell types. Note that MES cells still have the lowest ddhodge potential (earliest pseudotime). Also note the relatively similar values across GBM cell types. (G – I) Abstracted PhenoCycler maps of called cell types from the indicated sample. Note the close association of Intermediate cell types along the interface between MES-like and OPC-like cells. (J) Data from PhenoCycler indicating the percentage of cell type that is positive for Ki67 (proliferating) across both core and peripheral samples. Two-way ANOVA result: p = 0.035 for cell type and p = 0.012 for group. Tukey’s post-hoc MES-like vs OPC-like p = 0.044. (K) Single cell multiome data showing coverage plot of *NES* and *BCAN* ordered by inferred cellular pseudotime from RNA velocity. Note that as cells progress through pseudotime, *NES* gene expression (RNA) decreases while chromatin accessibility remains relatively similar.

**Video S1.** Animation demonstrating most probable path of MES cells over time, related to Figure 7.

**Video S2.** 3D plot with UMAP embeddings as the X and Y axes while the Z axis is the UMAP-based ddhodge potential. This produces a Waddington-like plot of cell states, related to Figure 7.

**Video S3**. 3D plot of combined GBM and developing brain cells. Axes are as in Video S2, related to Figure 7.

