## Supplementary figures and images for "Spatial epigenomic niches underlie glioblastoma cell state plasticity"

### Video S1

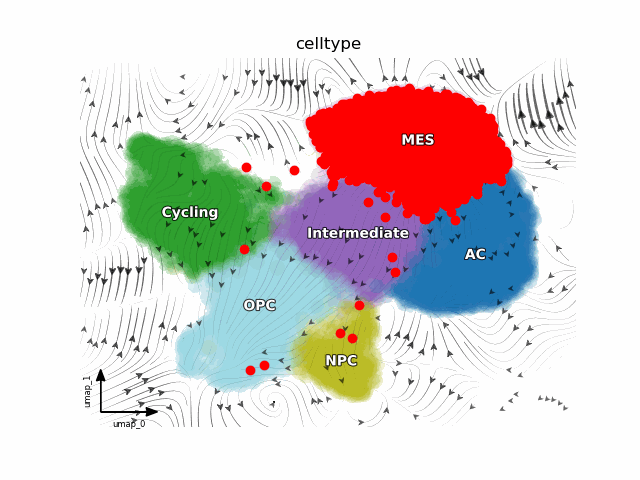

### Video S2

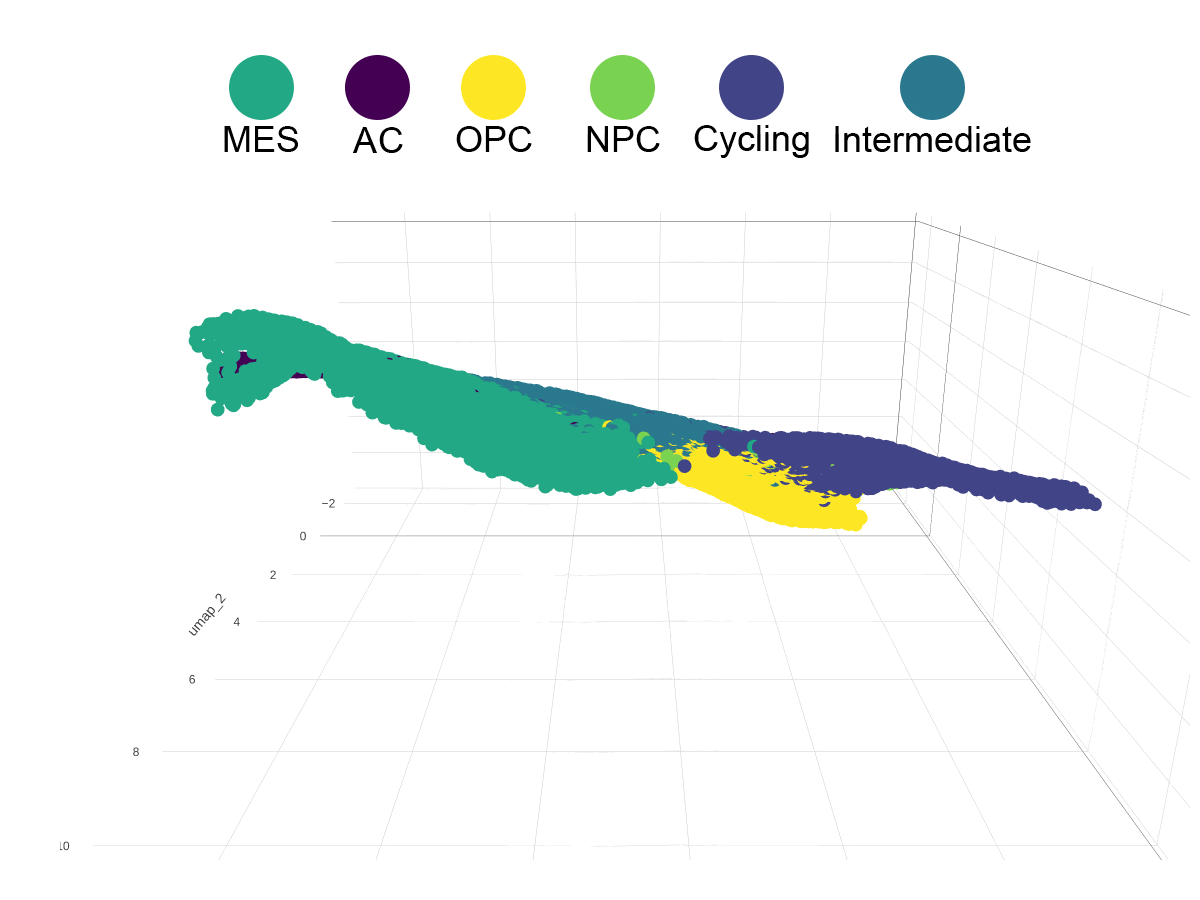

### Video S3

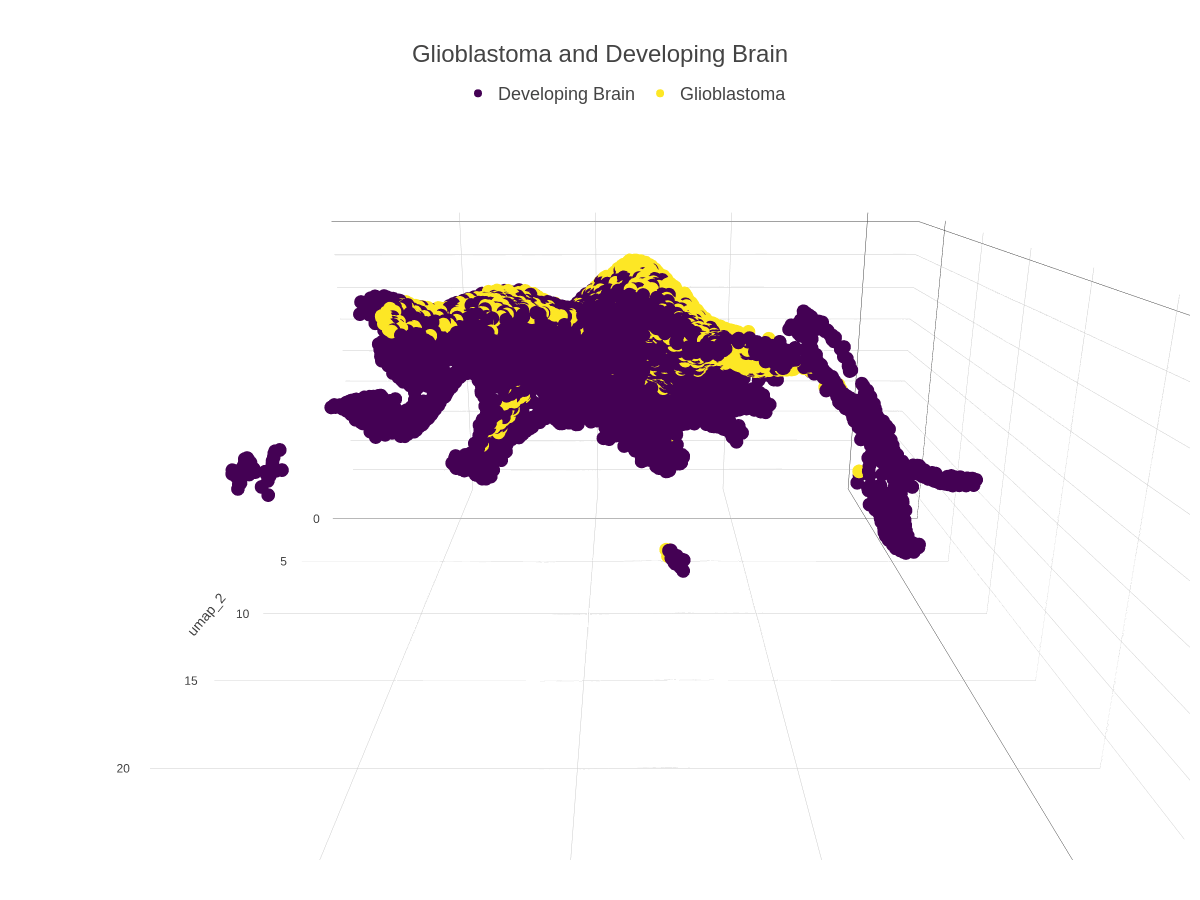
